# In colon cancer cells, fascin1 functions as a mechanosensor that transforms adherens junction mechanotransduction

**DOI:** 10.1101/2022.06.14.496198

**Authors:** Amin Esmaeilniakooshkghazi, Eric Pham, Sudeep P. George, Afzal Ahrorov, Fabian R. Villagomez, Michael Byington, Srijita Mukhopadhyay, Srinivas Patnaik, Jacinta C. Conrad, Monali Naik, Saathvika Ravi, Niall Tebbuttt, Jennifer Mooi, Camilla M Reehorst, John M. Mariadason, Seema Khurana

## Abstract

Fascin1 expression in colorectal carcinomas (CRCs) is linked to a clinically aggressive disease with poor prognosis. Despite that fascin1’s role in the etiology of CRCs has not been directly investigated. We show fascin1 expression in one-third of all CRCs underscoring the critical need to identify fascin1’s function in colorectal carcinogenesis. Here, we identify for the first time, fascin1’s role as a mechanosensor that modulates CRC cell adherens junction (AJ) plasticity to induce tumor growth and metastasis. We show that fascin1 expression drives protein sorting to transform AJ mechanotransduction and we reveal how these force-sensitive pathways activate oncogenic signaling in CRC cells. We made the novel finding that AJ remodeling by fascin1 also controls “collective plasticity” and bidirectional cell migration. Few studies have examined AJ plasticity in cancer cells, which remains poorly understood and has not been therapeutically targeted. Our findings could have widespread implications for understanding and treating metastatic carcinomas.

## Introduction

Fascin1 is an actin-bundling protein that is expressed throughout the gastrointestinal (GI) tissue in the mucosa and the muscle layers during mouse and human development.(De Arcangelis et al., 2004; Zhang et al., 2008) Notably, fascin1 expression in the developing GI tissue is intense in regions of ongoing differentiation, cell migration, and cell proliferation indicative of a role for fascin1 in cell-cell interactions, cell division, and cell migration.(De Arcangelis *et al*., 2004; Zhang *et al*., 2008) Nevertheless, fascin1 transcript and protein are absent from the normal adult intestinal epithelium.(Zhang *et al*., 2008) However fascin1 is re-expressed in colorectal carcinomas (CRCs), where its expression is correlated with a clinically aggressive disease with a higher incidence of metastasis and poor prognosis.(Hashimoto et al., 2006; Shi et al., 2020; Tan et al., 2013; Vignjevic et al., 2007) Moreover, in multivariate analysis, fascin1 has been identified as an independent factor for CRCs, and fascin1 positive CRCs exhibit a greater ability to invade lymph nodes, to develop extra-nodal tumor extensions, and have a higher likelihood of recurrence with a lower rate of disease-free and overall survival.(Hashimoto *et al*., 2006; Ristic et al., 2021; Tampakis et al., 2021) It is assumed that fascin1 drives colorectal tumor cell migration and invasion by assembling filopodia and invadopodia.(Lin et al., 2021; Liu et al., 2021) However, fascin1’s function in CRCs has not been directly investigated. There is emerging evidence from other cancer cells that fascin1 also has cell migration-independent functions linked to tumor cell proliferation, oncogenesis, metastatic colonization, anoikis resistance, chemoresistance, and cancer stemness although the molecular mechanism underlying these non-canonical functions of fascin1 remain unclear.(Barnawi et al., 2016; Li et al., 2014; Ristic *et al*., 2021) Consequently, the mechanism by which fascin1 promotes metastatic CRCs also remains unknown. The functional characterization of fascin1 could shed light on the biology driving metastatic CRCs.

During epithelial tissue development, repair and homeostasis, cell-cell adhesion junctions called adherens junctions (AJs) regulate initiation and stabilization of cell-cell adhesion as well as regulate the actin cytoskeleton, intracellular signaling and transcriptional regulation thereby ensuring tissue integrity while allowing for cell and tissue dynamics.(Fristrom, 1988) (Garcia et al., 2018) E-cadherin-β-catenin complex and other AJ associated proteins are mechanosensors and contractile forces generated by the actomyosin cytoskeleton initiate downstream signaling pathways that control tissue development by controlling cell fate including cell differentiation, cell proliferation, cell death and cell migration/invasion. Although AJs are stable in mature epithelia, the AJs between cells are dynamic during morphogenetic processes and in the adult organs during regenerative processes such as wound healing or during pathogenic processes such as cancer invasion.(Takeichi, 2014) While much is known about nascent AJ formation, remodeling of AJs in the mature epithelia is less well-understood. In tumor cells, AJs play an important role in survival, growth, invasion and metastasis.(Na et al., 2020; Padmanaban et al., 2019) Increased AJ plasticity is also related to the ability of cancer cells to move collectively which allows them to intravaste and extravasate more efficiently and makes them more resistant to cell death.(Aceto et al., 2014; Friedl et al., 2012; Grosse-Wilde et al., 2015; Joosse et al., 2015; Klymkowsky and Savagner, 2009; Revenu and Gilmour, 2009; Ribeiro and Paredes, 2014; Watanabe et al., 2014; Welch-Reardon et al., 2014; Yu et al., 2013; Zheng et al., 2017) Collective migration is also advantageous for tumor cells because, it eliminates the need for all cells to detect external signals for migration and it allows coupling of mechanical forces among cells for maximum plasticity.(Aceto *et al*., 2014; Friedl *et al*., 2012; Joosse *et al*., 2015; Klymkowsky and Savagner, 2009; Revenu and Gilmour, 2009; Ribeiro and Paredes, 2014; Watanabe *et al*., 2014; Welch-Reardon *et al*., 2014; Zheng *et al*., 2017) Compared to single cell, collective cell migration has several other advantages, allowing cells to respond more efficiently to external cues.(Theveneau et al., 2010) Consequently, tumor cells that maintain remodeled AJs are more aggressive, with higher metastatic risk and poorer clinical outcomes.(Bidard et al., 2014; Cristofanilli et al., 2004; Khoo et al., 2015; Singh and Settleman, 2010; Thiery et al., 2009; Yu *et al*., 2013) Despite the implications that pathological remodeling of AJ proteins is integral to transformation and tumor invasion, few studies have examined AJ remodeling in cancer cells.

In the normal crypt-villus axis migration of intestinal epithelial cells, moving cells remain fully attached to their neighbors and to the extracellular matrix as they move unidirectionally upwards as a cohesive sheet.(Nanba et al., 2015; Wong et al., 1998) While the mechanism guiding single cell migration are well understood, cell-intrinsic mechanisms that steer collectively migrating cells are not fully resolved. For instance, collective cell migration involves the integration of guidance cues between functionally and morphologically distinct cell populations (e.g. leader-follower) many of which have not been defined. During morphogenesis and during cancer invasion, another flexible type of collective cell migration occurs where some properties of single cell migration are acquired by collectively migrating cells, when sheets of cells and detached groups of cells maintain cell-cell adhesions while migrating as a loosely cohesive groups.(Haeger et al., 2014; Ilina et al., 2011; Osswald et al., 2015; Scarpa et al., 2015; Theveneau and Mayor, 2011) Such “collective plasticity” of tumor cells occurs when infiltrating local tissue.(Cheung and Ewald, 2016; Ilina and Friedl, 2009) Partial or hybrid epithelial-mesenchymal-transition (EMT) weakens cell-cell junctions while retaining E-cadherin expression which contributes to tumor invasion of loosely connected cancer cells.(Jolly et al., 2015; Nieto et al., 2016) However, the mechanisms regulating such “collective plasticity” of cancer cells remain unclear. Moreover, the proteins regulating the remodeling and weakening of cell-cell junctions remain unidentified.

*In vivo* studies have shown that imposing uniaxial deformation to *Drosophila* embryos is sufficient to induce β-catenin nuclear localization and ectopic expression of Wnt target genes (e.g. Twist).(Farge, 2003) A similar pathway has also been described for nuclear accumulation of β-catenin in gastrulating zebrafish embryos.(Brunet et al., 2013) Consistent with that, long-term mechanical pressure exerted by colon tumor growth has been shown to trigger proliferation of surrounding non-tumorigenic cells *via* Wnt/β-catenin activation. (Fernandez-Sanchez et al., 2015) Additionally, Frizzled, a well-known Wnt receptor also responds to mechanical stimulation resulting in Wnt/β-catenin activation.(Rotherham and El Haj, 2015) These findings point to a conserved role for mechanosensitive Wnt/β-catenin signaling during morphogenesis and tumorigenesis. Mechanistically, it has been proposed that actomyosin activity at AJs mediates the effects of mechanical forces on Wnt/β-catenin activation.(Hall et al., 2019; Hirata et al., 2017) However, few studies have investigated how force-sensitive pathways are regulated in cancer cells to regulate oncogenic Wnt/β-catenin signaling.

Using a large cohort of CRC patients, we show that fascin1 is upregulated in one-third of all CRCs. Our study also identifies a correlation between high fascin1 expression and more poorly differentiated/high grade CRCs, which grow and metastasize more rapidly. These findings underscore the critical need to understand the function of fascin1 in colon carcinogenesis. In this study we identify a novel function of fascin1 in CRC cell AJ remodeling that influences both tumor growth and invasion. We show for the first time, that in CRC cells fascin1 reorganizes AJ-associated actin cytoskeleton to assemble dynamic cell-cell adhesions including the assembly of ultra-long and large tumor microtube-like intercellular junctions that modulate the oncogenic behavior of CRC cells. We demonstrate that by remodeling AJ-associated actin cytoskeleton fascin1 directly activates mechanosensitive Wnt/β-catenin signaling. Additionally, we demonstrate that fascin1’s AJ remodeling property induces “collective plasticity” allowing cells to migrate as a loosely cohesive group of cells with different migration speeds and patterns. We show for the first time, the direct role of fascin1 in generating bidirectional epithelial cell migration. This is, to the best of our knowledge, the first identification of an actin-binding protein that regulates cell-intrinsic reverse and bidirectional cell migration. By overexpressing fascin1 in non-transformed epithelial cells, we identify fascin1’s direct role in AJ remodeling that contributes to cellular plasticity programs that influence tumor initiation and progression. Fascin1 is expressed in a wide range of carcinomas where its expression has also been correlated with a clinically aggressive disease associated with high mortality.(Tan *et al*., 2013) Based on that we propose that our findings could have wider implications for understanding the molecular mechanisms driving AJ plasticity in carcinogenesis and this information could be leveraged to develop novel therapies to treat metastatic carcinomas.

## Results

### Fascin1 is upregulated in a significantly large number of CRCs and its expression correlates with more aggressive tumors

Varying rates of fascin1 expression in CRCs have been reported consequently, there is no clear consensus on the frequency of fascin1 expression in CRCs. This is in large part because many of these studies have examined fascin1 expression at different stages of tumorigenesis. To characterize fascin1’s role in colorectal carcinogenesis, we elected first to analyze fascin1 expression in a large cohort of 299 stage IV CRCs, from patients who participated in the Phase III MAX clinical trial.(Tebbutt et al., 2010) Analysis of fascin1 expression using the Hashimoto *et al* scoring criteria revealed low to high fascin1 expression in 34.4% of cases (score 1-3) while 22.4% of cases expressed moderate to high fascin1 protein levels (score 2-3; Fig 1A).(Hashimoto *et al*., 2006) Additionally, in this patient cohort we found that fascin1 was more highly expressed in high grade/poorly differentiated CRCs (G1>G2>G3; Fig 1B) confirming fascin1’s association with more aggressive and invasive colorectal tumors. Understanding the underlying molecular mechanism of fascin1 could improve treatment and increase survival for at least a third of all CRC patients.

**Figure 1.**
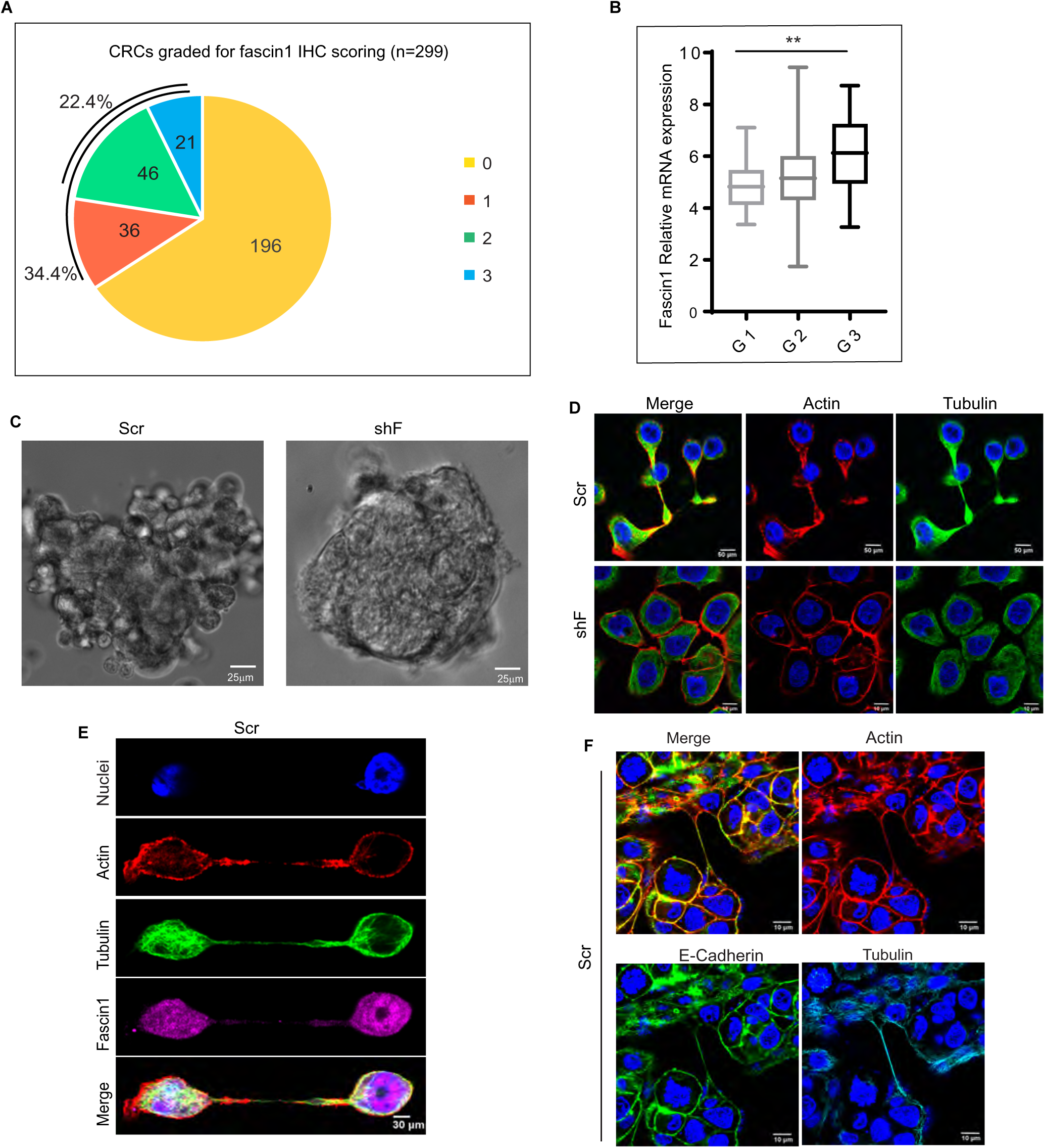
The role of fascin1 in CRC. **(A)** CRCs were graded for fascin1 protein expression and pie chart show the correlation of fascin1 with CRCs. **(B)** Fascin1expression increases significantly with increasing tumor grade (G1>G2>G3, **p<0.01, n=270). **(C)** Phase contrast image of HT-29/19A-Scr and shF cells grown as suspension 48 h tumor spheroids. Scale bar, 25 µm. **(D)** Confocal images of sub confluent HT-29/19A-Scr and HT-29/19A-shF cells show distribution of actin (red) and tubulin. Nuclei were counter stained with DAPI. Scale bar, 50 µm (Scr) and 10 µm (shF). **(E)** Confocal image of sub confluent HT-29/19A-Scr cells shows distribution of actin (magenta), tubulin (red), and fascin1 (green). Nuclei were counter stained with DAPI. Scale bar, 30 µm. **(F)** Confocal image of HT-29/19A-Scr cells shows distribution of actin (red), tubulin (cyan) and E-cadherin (green). Nuclei were counter stained with DAPI. Scale, 10 µm.

### In CRC cells fascin1 remodels adherens junctions and forms ultra-long and large tumor microtube-like intercellular connections

To identify the function of fascin1 in the pathogenesis of colorectal cancer we performed loss-of-function studies by knockdown (KD) of fascin1 in HT-29/19A CRC cells. Multiple fascin1 targeting shRNAs were used all of which markedly reduced endogenous fascin1 protein expression in the parental cell line but did they did not affect shRNA-refractive cDNA expression of EGFP-Fascin1 confirming on-target specificity of the shRNA constructs (Supplementary Fig 1A).(Xu et al., 2009) Spheroids of HT-29/19A cells expressing scrambled shRNA (HT-29/19A-Scr) grew as loosely aggregated clusters of cells with a ‘grape-like morphology.’ Comparatively, fascin1 KD cells (HT-29/19A-shF) formed tightly organized, compact spheres (Fig 1C), implicating a role for fascin1 in the remodeling of cell-cell adhesions. In sub-confluent cell cultures, HT-29/19A-Scr cells formed nascent cell-cell adhesions that appeared as long filopodia-like protrusions that morphed into lamellipodia-like structures at the distal ends as two cells established cell-cell contact (Fig 1D). In contrast, HT-29/19A-shF cells extended much smaller lamellipodial protrusions as two cells approached each other, similar to nascent adherens junction (nAJ) assembly by normal intestinal epithelial cells.(Baum and Georgiou, 2011) Notably, Scr cells formed these protrusions even in the absence of direct cell-cell contact highlighting fascin1’s autonomous role in generating these large cell surface protrusions (Supplementary Fig 1B). Alternatively, Scr cells formed long intercellular conduits that lacked lamellipodia-like distal ends (Fig 1E). These remodeled cell-cell junctions contained fascin1, F-actin and tubulin (Figs 1D-1E) as well as E-cadherin indicating these protrusions are indeed nAJs (Fig 1F). In these nAJs, fascin1 localized to both the filopodia-like and lamellipodia-like protrusions. By measuring the rate of assembly, we determined that fascin1, F-actin and tubulin are required to assemble these nAJs (Supplementary Figs 1C-1D). In normal intestinal epithelial cells, AJs are assembled by a symmetrical ‘push and pull’ mechanism.(Krendel et al., 1999) Remarkably, fascin1 expressing Scr cells assembled AJs asymmetrically, where only one cell extended the protrusion (Cell 1, Fig 2A) with a lamellipodia-like distal end (red arrowhead) or long intercellular conduits (cyan arrowhead) while the neighboring cell assembled actin bundles that appear like stress fibers that attach to the E-cadherin and β-catenin foci (white arrowheads). In contrast, shF cells assembled AJs symmetrically where both cells generated similar protrusions and displayed similar actin organization at the cell-cell contacts. To determine how fascin1 expression could modulate normal epithelial cell AJ assembly and to identify fascin1’s function independent of epithelial transformation, we performed gain-of-function studies by overexpressing EGFP tagged fascin1 in the normal epithelial MDCK cells (Supplementary Fig 1E). Remarkably, fascin1 expression in normal epithelial cells likewise remodeled cell-cell adhesions resulting in the assembly of ultra-long nAJs similar to those seen in CRC cells (Figs 2B). Notably, some MDCK EGFP-Fascin1 cells also initiated nAJ assembly by forming protrusions that resembled the sprouting phenotype of endothelial cells undergoing anastomosis (Fig 2C); or had a growth cone-like morphology (Fig 2D) reminiscent of fascin1’s function in these tissue.(Dent et al., 2011; Ma et al., 2013) More remarkably, these ultra-long nAJs were large enough to encompass functional organelles including mitochondria and lysosomes (Fig 2E).

**Figure 2.**
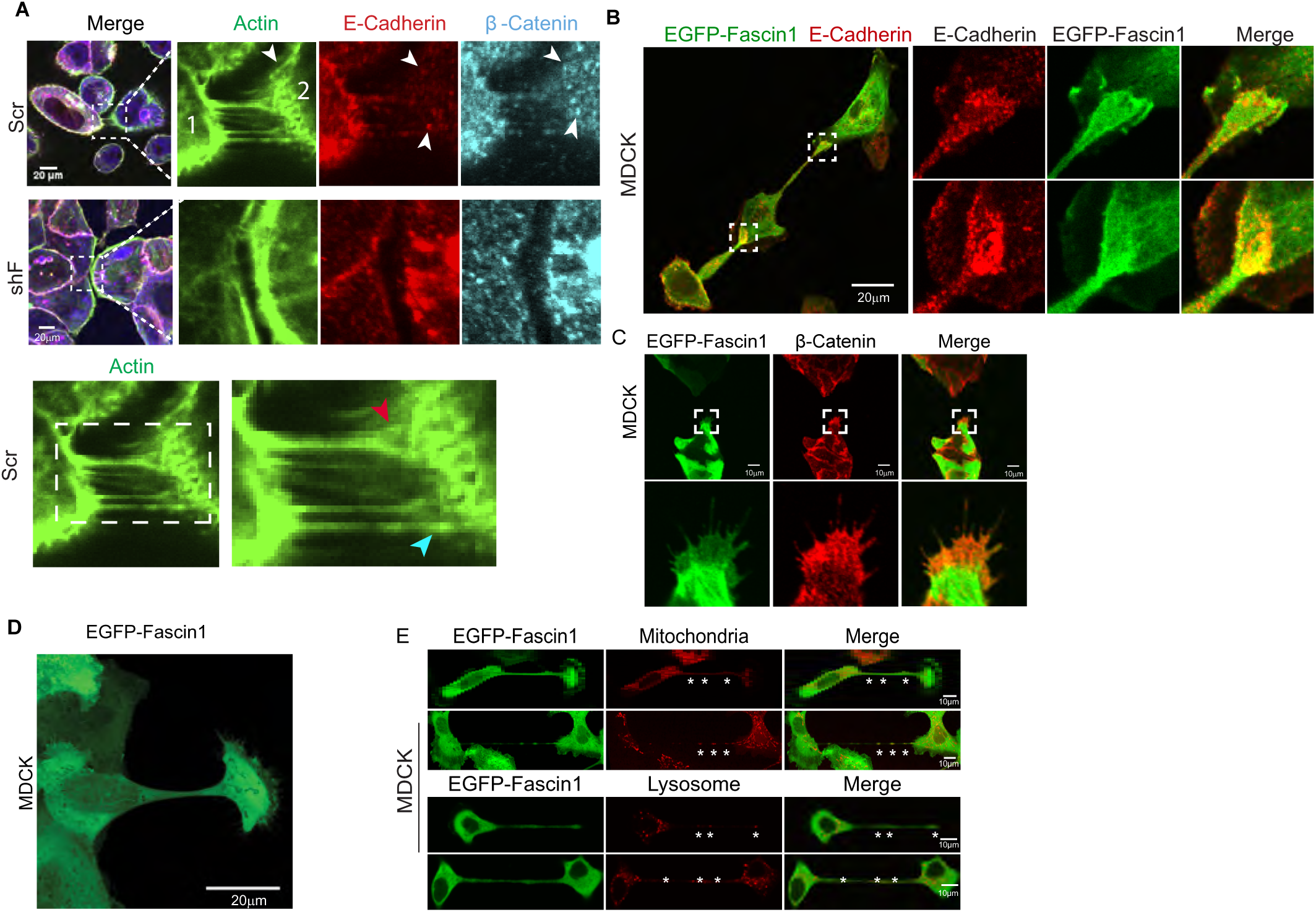
In CRC cells fascin1 remodels AJ assembly. **(A)** Confocal image of Scr and shF cells shows distribution of actin (green), E-cadherin (red) and β-catenin (cyan) as two cells (cell 1 and 2) assemble nAJs. Arrowheads identify, oblique actin bundles, E-cadherin and β-catenin foci (white). Elongated nascent cell-cell junctions with or without distal lamellipodia (cyan). Right panel is a magnification of the boxed area. Scale, 20 µm. **(B)** Confocal image of MDCK EGFP-Fascin1 cells shows distribution of fascin1 (green) and E-cadherin (red). Scale bar, 20 µm. **(C)** Confocal image of MDCK EGFP-Fascin1 cells shows the distribution of fascin1 (green) and β-catenin (red) during nAJ assembly. Bottom panels are magnification of the boxed areas shown above. Scale bar, 10 µm. **(D)** Confocal image MDCK EGFP-Fascin1 cells shows growth-cone like cellular protrusion during nAJ assembly. Scale bar, 20 µm. **(E)** Live confocal immunofluorescence of MDCK EGFP-Fascin1 cells labeled with MitoTracker-Red and LysoTracker Red show the presence of functional mitochondria and lysosomes in these elongated nascent AJs. Asterisks highlight mitochondria and lysosomes, respectively. Scale bar, 10 µm.

To characterize the effect of fascin1 expression on mature epithelial AJs, we examined AJs in confluent MDCK cells expressing EGFP-Fascin1. Remarkably, fascin1 also remodeled mature AJs forming ultra-long cell-cell junctions in confluent cell cultures that connected two non-adjacent and significantly distant cells (Fig 3A, cell 1 is connected to cell 6). These remodeled AJs hovered over the confluent monolayer, which facilitated two distant cells to connect in this manner (Fig 3B). To analyze how fascin1 remodels ‘mature, established’ epithelial AJs, small clusters of MDCK EGFP and MDCK EGFP-Fascin1 cells were stimulated with hepatocyte growth factor (HGF) at concentrations that do not induce epithelial cell scatter. While the MDCK EGFP cells maintained stable cell-cell adhesions with few short-lived changes, similar to normal epithelial (Supplementary Video 1, Fig 3C), the MDCK EGFP-Fascin 1 cells continuously remodeled their cell-cell adhesions, dissolving and reforming them resulting in loosely connected cell clusters (Supplementary Video 2, Fig 3C). These dynamically remodeled cell-cell adhesions differed in length and lifespan which were positively correlated (Fig 3D). Morphologically similar structures called tunneling nanotubes (TNTs) are short-lived intercellular bridges with lifespans of less than 60 min while tumor microtubes (TMs) have lifespans greater than 60 min.(Osswald *et al*., 2015; Rustom et al., 2004) Based on the morphological characteristics of these remodeled intercellular junctions and their lifespan, we refer to these intercellular connections assembled by fascin1 as tumor microtube-like (TM-like). Together, these data demonstrate that fascin1 expression in epithelial cells including CRC cells remodels AJ assembly as well as the stability of mature AJs.

**Figure 3:**
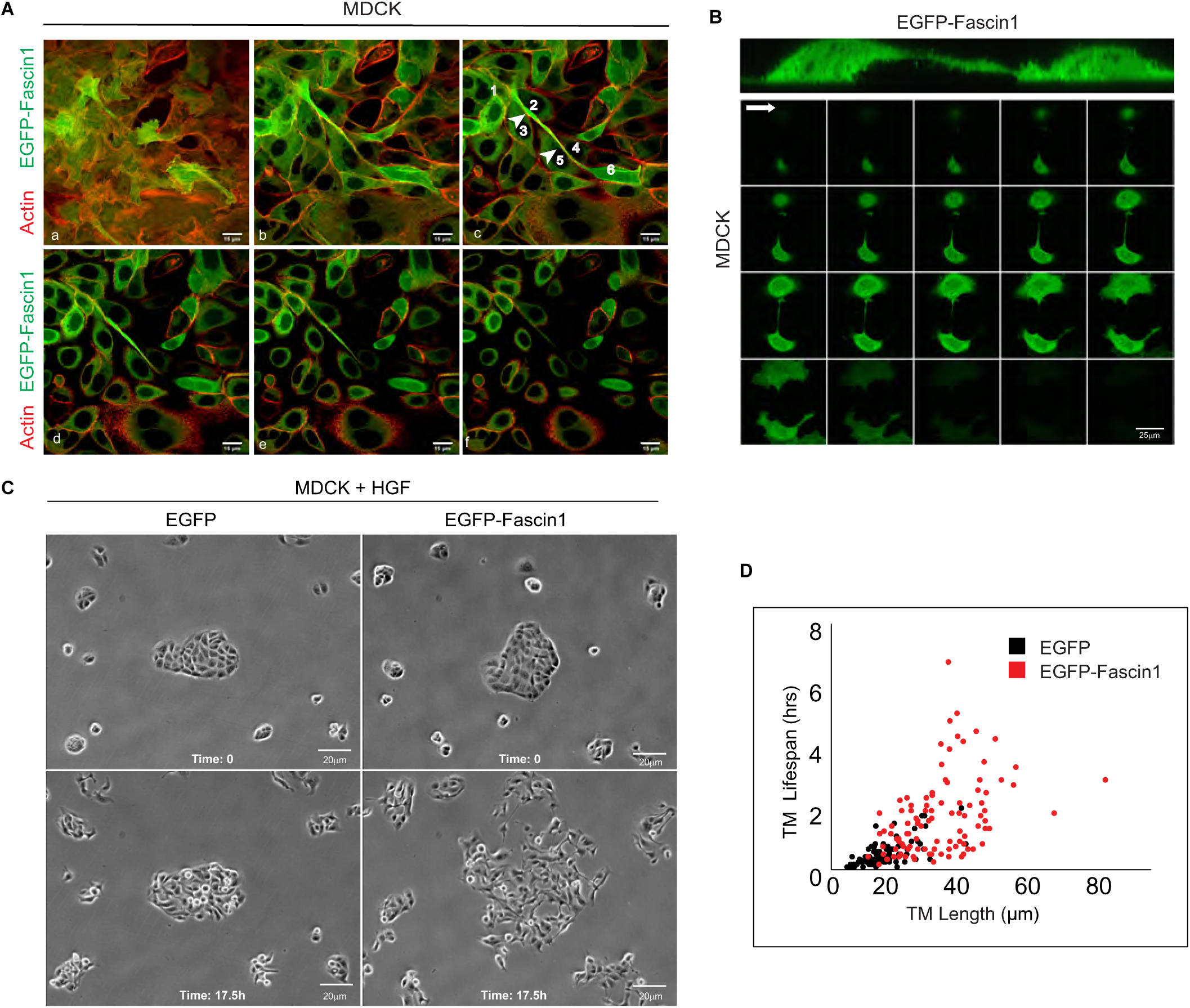
Fascin1 remodels mature AJs and assembles ultra-long, large and long-lived tumor microtube-like intercellular connections. **(A)** Confocal immunofluorescence z-stacks (xy images;1 µm each from (a) to (f)) show the distribution of fascin1 (green) and actin (red) in confluent MDCK EGFP-Fascin1 cells. Arrowhead identifies TM-like intercellular junctions connecting cell 1 to cell 6. Scale bar, 15 μm. **(B)** Live confocal immunofluorescence z stacks (xy images; 1 μm) of MDCK EGFP-Fascin1 cells show TM-like cell-cell junction suspended over the confluent monolayer. Scale bar, 25 μm. **(C)** Phase contrast images show clusters of MDCK EGFP and MDCK EGFP-Fascin1 cells at time 0 and at 17.5 h post-treatment with HGF. Scale bar, 20 μm. **(D)** Correlation of TM-like remodeled cell-cell junction length (μm) and lifespan (hrs) in HGF treated MDCK EGFP and MDCK EGFP-Fascin1.

### Fascin1 transforms adherens junction mechanotransduction in CRC cells

In epithelial cells, the actin cytoskeleton is important for the stabilization of AJs, but it is also the force-generating mechanism required for AJ remodeling e.g. during wound healing and the actin cytoskeleton regulates AJ mechanotransduction.(Hoffman and Yap, 2015; Ladoux et al., 2015; Michael and Yap, 2013; Schnittler et al., 2014) In normal intestinal epithelial cells, mature AJs are stable, continuous, and associated with actin bundles running parallel to the plasma membrane that appear at cell-cell margins overlapping with the cortical actin.(Yonemura, 2011) However, in fascin1 expressing CRC cells, we identified AJs with actin bundles arranged perpendicular to the cell-cell junctions that appear as intercellular stress fibers forming end-on attachments with the discontinuous E-cadherin foci (Fig 4A, Top). In stark contrast, shF CRC cells formed continuous AJs with F-actin and E-cadherin distribution overlapping with the cell-cell margins similar to normal intestinal epithelial cells. There was no discernable difference in the lateral junctions between Scr and shF cells (Fig 4A, Middle). Gain-of-function studies were performed in a CRC cell line that lacks endogenous fascin1 protein by overexpressing EGFP-Fascin1 in HT-29 cells (Supplementary Figs 2A-2B). Overexpression of fascin1 in HT-29 cells resulted in remodeling of AJ associated actin cytoskeleton, confirming fascin1’s direct role in this modification. This function of fascin1 was also recapitulated in the non-transformed epithelial MDCK cells overexpressing EGFP-Fascin1 (Supplementary Fig 2C); and in the rescue experiments performed in the fascin1 KD shF cells by overexpressing EGFP-Fascin1 (Supplementary Fig 2D). Together, these data demonstrate that fascin1’s function in AJ remodeling is not cell type-specific and is independent of oncogenic transformation implying that fascin1 re-expression in the intestinal epithelium generates this phenotype irrespective of other oncogenic signals.

**Figure 4.**
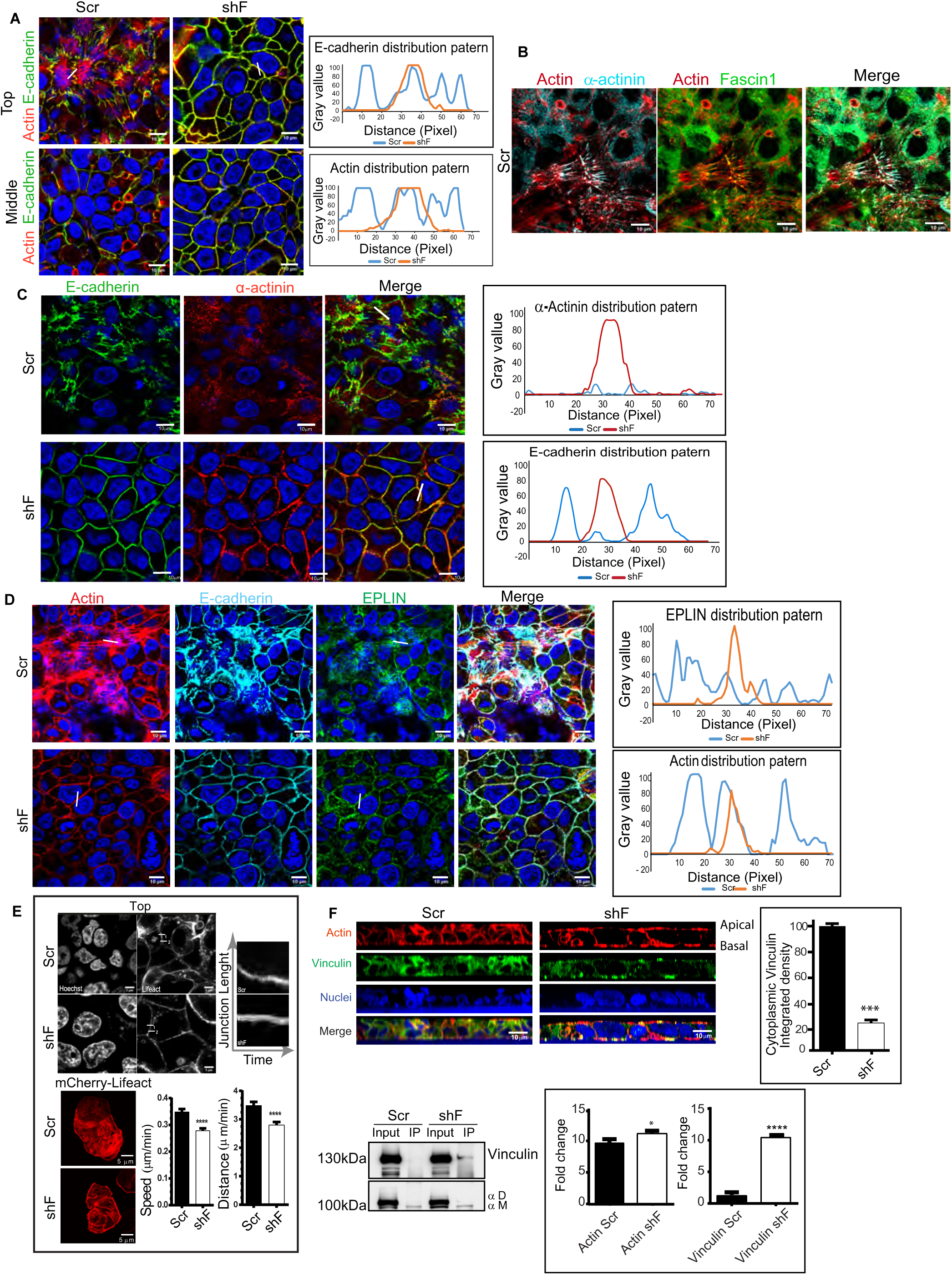
Fascin1 remodels AJ mechanotransduction in CRC cells. **(A)** Confocal microscopy shows the distribution of actin (red) and E-cadherin (green) at the AJs (Top) and lateral junctions (Middle) of HT-29/19A-Scr and shF cells. Nuclei were counter stained with DAPI. Plot profiles were generated by measuring the distribution of actin and E-cadherin in ROI highlighted by a white line, 10 μm long. Scale, 10 μm. **(B)** Virtual Channel confocal microscopy shows the AJ associated actin (red), fascin1 (green) and α-actinin (cyan) in Scr cells. Scale bar, 10 μm. **(C)** Confocal immunofluorescence of HT-29/19A-Scr and shF cells show the distribution of E-cadherin (green) and α-actinin (red) at the AJs. Nuclei were counter stained with DAPI. Plot profiles were generated by measuring the distribution of E-cadherin and α-actinin in ROI highlighted by a white line, 10 μm long. Scale bar, 10 μm. **(D)** Virtual Channel confocal immunofluorescence of HT-29/19A-Scr and shF cells shows distribution of actin (red), E-cadherin (cyan), and EPLIN (green) at the AJs. Nuclei were counter stained with DAPI. Plot profiles were generated by measuring the distribution of EPLIN and actin in ROI highlighted by a white line, 10 μm long. Scale bar, 10 μm. **(E)** Live cell imaging of HT-29/19A-Scr and shF cells stably expressing mCherry-Lifeact. Nuclei were counter stained with Hoechst. Scale bar, 5μm. Kymograph analysis of mCherry-Lifeact at AJs of Scr and shF cells measured at two ROI (labeled 1 and 2, 10 μm each) shows difference in speed (μm/min) of migration and distance (μm/min) moved by AJ-associated F-actin in Scr and shF cells. Asterisk (****) denotes p<0.001, n=6. **(F)** Confocal images of Scr and shF cells show xz distribution of vinculin (green) and actin (red). Nuclei were counter stained with DAPI. Scale bar, 10 μm. Accumulation of cytoplasmic vinculin in Scr and shF cells was measured from three independent experiments (***) denotes p<0.001, n=3. Scr and shF cell lysates were immunoprecipitated with α-catenin antibody and immunoblotted with vinculin antibodies. Asterisks (*) denote as follows: *** denotes p<0.001, n=3 and **** p<0.0001, n=3.

*In vitro* fascin1 and α-actinin mutually exclude each other to form discrete fascin1 or α-actinin cross-linked actin networks.(Winkelman et al., 2016) We reasoned that in CRC cells, fascin1 re-expression could result in a similar competitive exclusion of AJ associated actin-bundling proteins α-actinin and potentially EPLIN which could explain fascin1’s AJ remodeling property in CRC cells. If true, such changes would also have significant implications for AJ mechanotransduction, because both α-actinin and EPLIN regulate AJ mechanosensitivity.(Le et al., 2017; Taguchi et al., 2011) Indeed, in CRC Scr cells, fascin1 and α-actinin segregate to different areas of the AJ-associated actin cytoskeleton (Fig 4B). While α-actinin co-localized with the oblique contractile actin bundles of the AJs, fascin1 was associated with the actin bundles where they attach to the E-cadherin foci (Fig 4B). This close proximity of fascin1 to the E-cadherin foci also suggested to us that in CRC cells, fascin1 could play a direct role in modulating AJ mechanotransduction. As expected, in Scr cells we identified a competitive exclusion of α-actinin and EPLIN away from the AJs to the cytoplasm (Figs 4C-4D). In shF cells, the α-actinin and EPLIN distribution was restored to the cell-cell margins and the AJs similar to normal intestinal epithelial cells. Fascin1 also displaced α-actinin away from focal adhesions (FAs) of Scr cells (Supplementary Fig 2E). There was no significant difference in total α-actinin protein levels between control and fascin1 KD HT-29/19A cells (Supplementary Fig 2E). To quantitatively measure the effects of fascin1 on AJ actin dynamics, we transfected HT-29/19A Scr and shF cells with mCherry-Lifeact (Fig 4E). As anticipated, kymographs revealed greater movement and greater velocity of movement of F-actin at the AJs of Scr cells compared to shF cells indicative of greater AJ plasticity. In normal epithelial cells, α-catenin is a key mechanosensor and monomeric α-catenin (α^M^) couples the E-cadherin-β-catenin complex to the actin cytoskeleton and recruits vinculin to reinforce the AJ stability while the α-catenin that dissociates from the cadherin-catenin complex homodimerizes in the cytoplasm (α^D^).(Desai et al., 2013; Seddiki et al., 2018; Yao et al., 2014) In this manner α-catenin and vinculin function as major mechanosensors that transmit the force of actomyosin contraction to the E-cadherin-β-catenin complex.(Fernandez-Sanchez *et al*., 2015; Przybyla et al., 2016; Yonemura et al., 2010) To directly measure the effect of fascin1 KD on AJ mechanotransduction, we evaluated the association of vinculin with the AJs and the association of vinculin with α-catenin.(Seddiki *et al*., 2018) While significant displacement of vinculin away from AJs (apical) and FAs (basal) to the cytoplasm was observed in HT-29/19A-Scr cells, fascin1 KD in HT-29/19A-shF cells restored vinculin localization to these cell adhesion sites similar to vinculin’s distribution in normal epithelial cells (Fig 4F; Supplementary Fig 2E).(le Duc et al., 2010) There was no significant difference in total vinculin protein levels between Scr and shF cells (Supplementary Fig 2E). Quantitative co-immunoprecipitation (IP) assay with an antibody that specifically immunoprecipitates monomeric α-catenin revealed that fascin1 knockdown also resulted in higher levels of vinculin in complex with α-catenin (Fig 4F). Together these data show that by simply competing with other actin-bundling proteins fascin1 reorganizes cellular actin cytoskeleton and remodels the AJs by displacing key AJ mechanosensors. Previously fascin1 has been shown to play a role in stress fiber organization at the focal adhesions.(Elkhatib et al., 2014) Our findings here that fascin1 displaces α-actinin and vinculin from FAs, also provides a molecular mechanism for fascin1’s function in FA remodeling.

### Fascin1 activates mechanosensitive Wnt/β−catenin signaling

Wnt activation in response to changes in AJ mechanotransduction plays an important role during morphogenesis and in tumorigenesis when these conserved embryonic mechanosensitive pathways are reactivated in the adult tissue.(Fernandez-Sanchez *et al*., 2015; Whitehead et al., 2008) We reasoned that by remodeling CRC cell AJ mechanotransduction, fascin1 could drive oncogenic Wnt activation. Indeed, subcellular fractionation showed that fascin1 KD significantly inhibited the nuclear accumulation of β-catenin (60.2%, n=3, p<0.001), and caused a redistribution of β-catenin from the nucleus and cytoplasm to the cell-cell margins and the AJs without impacting total β-catenin or E-cadherin protein levels (Fig 5A, Supplementary Figs 3A). Most notably, re-expression of EGFP-Fascin1 in the fascin1 KD shF cells resulted in the redistribution of β-catenin from the cell-cell margins and AJs back to the nucleus (Fig 5A) validating fascin1’s direct role in nuclear accumulation of β-catenin. Actively invading Scr cell tumor spheroids confirmed significant nuclear accumulation of β-catenin (Fig 5B). Nuclear accumulation of β-catenin was also identified in non-transformed MDCK cells overexpressing EGFP-Fascin1 further validating the direct role of AJ mechanotransduction in fascin1-mediated Wnt activation (Supplementary Fig 3B). Likewise in the CRC patient cohort high fascin1 protein expression (score 2-3) was associated with widespread (score 2) accumulation of nuclear β-catenin validating the pathophysiological significance of this function of fascin1 (Fig 5C). Fascin1 KD in CRC cells also resulted in reduced expression of Wnt target genes including the marker for stemness, *Cd44,* the ECM remodeling proteins, matrix metalloproteinase, *Mmp7* and *Mmp9* and proto-oncogene *c-Myc* which inhibits differentiation, promotes cell-cycle progression, promotes self-renewal, and chemoresistance including anti-EGFR resistance in colon cancer cells (Fig 5D).(Strippoli et al., 2020; Zhang et al., 2019) Reduced Wnt activation was confirmed by chromatin immunoprecipitation (ChIP-PCR) assay with the β-catenin antibody (Supplementary Fig 3C). Fascin1’s function as a driver of Wnt activation was validated with the Super8x TOP-Flash luciferase TCF-LEF reporter assay (Supplementary Fig 3D). The reduction of Wnt activation induced by fascin1 KD was also accompanied by significant changes in cell morphology and oncogenic behavior of cells such as reduced anchorage-independent growth (Fig 5E), restoration of tight junction morphology (Fig 5F), and restoration of contact inhibition (Fig 5F). Loss of contact inhibition was phenocopied in the rescue experiments performed in fascin1 KD shF cells overexpressing EGFP-Fascin1 cells (Supplementary Fig 4A) and in the gain-of-function studies performed in MDCK cells overexpressing EGFP-Fascin1 (Supplementary Fig 4B). Additionally, fascin1 KD in other CRC cell lines e.g. HCT-116 recapitulated fascin1’s function in loss of contact inhibition, suggesting a conserved function for fascin1 in all CRC cells (Supplementary Figs 4C). In all these cell types, fascin1 expression resulted in multi-layering in contrast to the KD cells that grew as monolayers. Together, our studies demonstrate that by activating Wnt target genes fascin1 plays a direct role in enhancing oncogenic behavior of epithelial cells.

**Figure 5.**
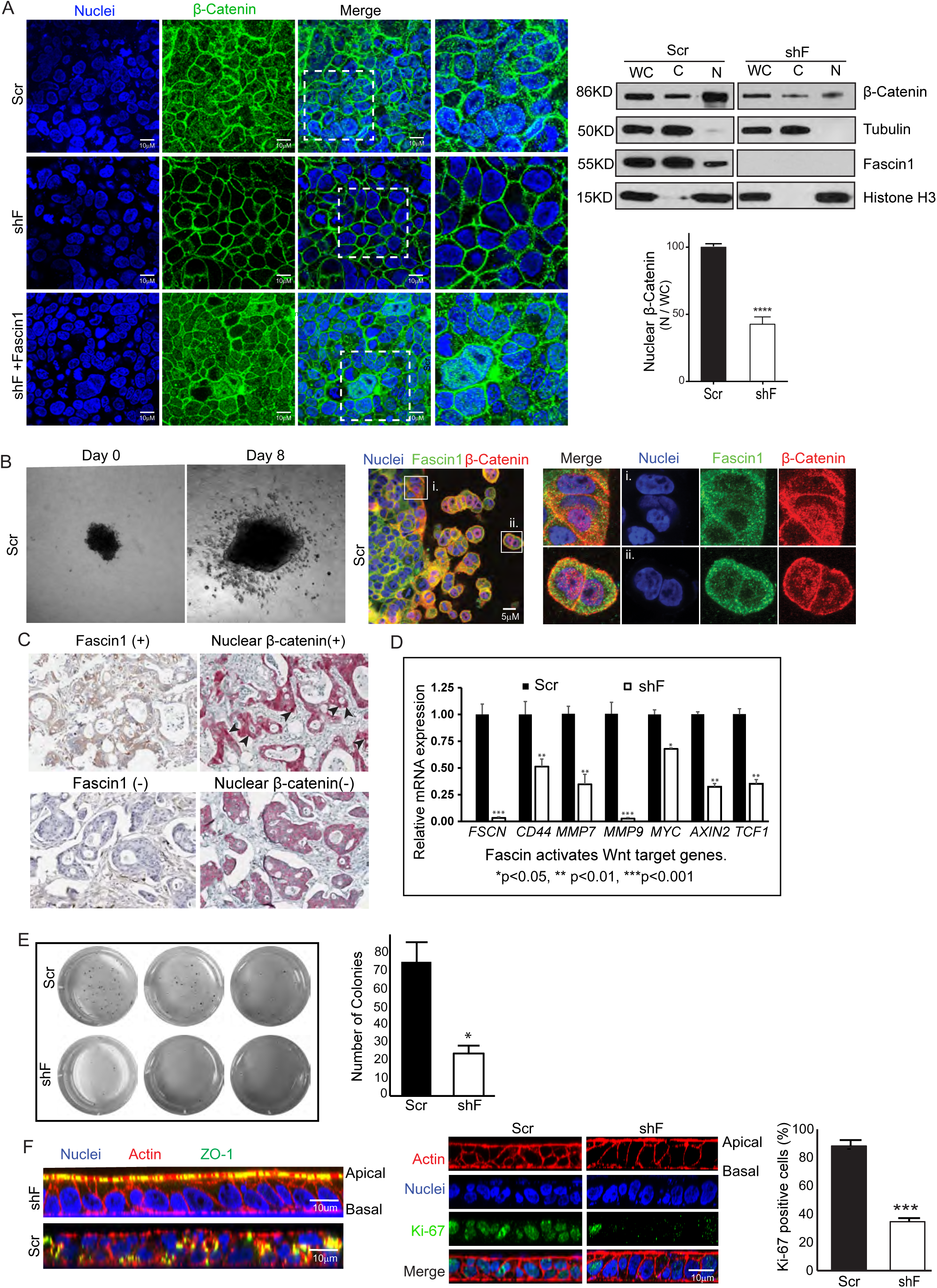
Fascin1 activates mechanosensitive Wnt/β-catenin signaling in CRC cells. **(A)** Confocal immunofluorescence of HT-29/19A-Scr, shF and shF expressing Fascin1 shows the distribution of total β-catenin (green). Nuclei were counter stained with DAPI. Scale bar, 10 μm. Subcellular fractionation assays were performed to quantify the distribution of β-catenin in the nuclear (N), cytoplasmic (C) and whole cell (WC) lysate fractions. Asterisk (****) denotes p<0.0001, n=3. **(B)** Phase contrast image shows actively invading Scr cell tumor spheroids at day 0 and at the end of day 8. Confocal images show the distribution of fascin1 (green) and β-catenin (red) in actively invading Scr cell tumor spheroids. Nuclei were counter stained with DAPI. Right panels (i) and (ii) are magnifications of the boxed areas labelled (i) and (ii). **(C)** Immunohistochemistry in paired panels show fascin1 and widespread nuclear β-catenin accumulation in the same colorectal tumor. Arrowheads highlight widespread accumulation of nuclear β-catenin in high fascin1 expressing tumors. **(D)** Quantitative RT-PCR for Wnt target genes using total cellular mRNA extracted from Scr and shF cells. Asterisks (*) denote as follows: * p<0.05, ** p<0.01, *** p<0.001 (n=6). **(E)** Phase contrast image of anchorage-independent colonies of HT-29/19A-Scr and shF cells. Asterisk (*) denotes p<0.05, n=3. **(F)** Confocal immunofluorescence shows xz distribution of ZO-1 (green) and actin (red) (left panels) and xz distribution of actin (red) and Ki67 (green; right panels) in Scr and shF cells. Nuclei were counter stained with DAPI. Asterisk (***) denotes p<0.001, n=3.

### Fascin1 activates mechanosensitive Wnt/β-catenin signaling to accelerate primary and secondary tumor growth

We reasoned that by activating Wnt target genes that promote cell proliferation and cell migration/invasion fascin1 could accelerate colorectal tumorigenesis. To test this idea, we assessed the effects of fascin1 KD on primary and secondary tumor growth of CRC cells *in vivo*. Fascin1 KD significantly reduced primary tumor weight compared to control cells (Fig 6A; p<0.01, n=6) and reduced the number of actively proliferating tumor cells validating fascin1’s role in the etiology of colorectal tumor growth (Supplementary Fig 5A). More importantly, fascin1 KD induced a re-localization of total and active (NP) β-catenin from the nucleus and cytoplasm to the cell membrane (Figs 6B-6C, respectively). We used the active (NP) β-catenin antibody to demonstrate the presence of active cytoplasmic β-catenin in these tumors. Most remarkably, we identified the ultra-long TM-like cell-cell junctions within the primary tumors in mice (Fig 6D). Fascin1 KD significantly reduced secondary tumor growth in mice (Fig 6E; p<0.01, n=8) and prevented the cytoplasmic and nuclear accumulation of total and active β-catenin (Fig 6E). Fascin1 KD also reduced CRC cell metastasis in a zebrafish embryo metastasis assay (Supplementary Fig 5B) validating this as a conserved function of fascin1. These data demonstrate that mechanosensitive Wnt activation regulated by fascin1 accelerates and augments colorectal tumor growth and metastasis.

**Figure 6.**
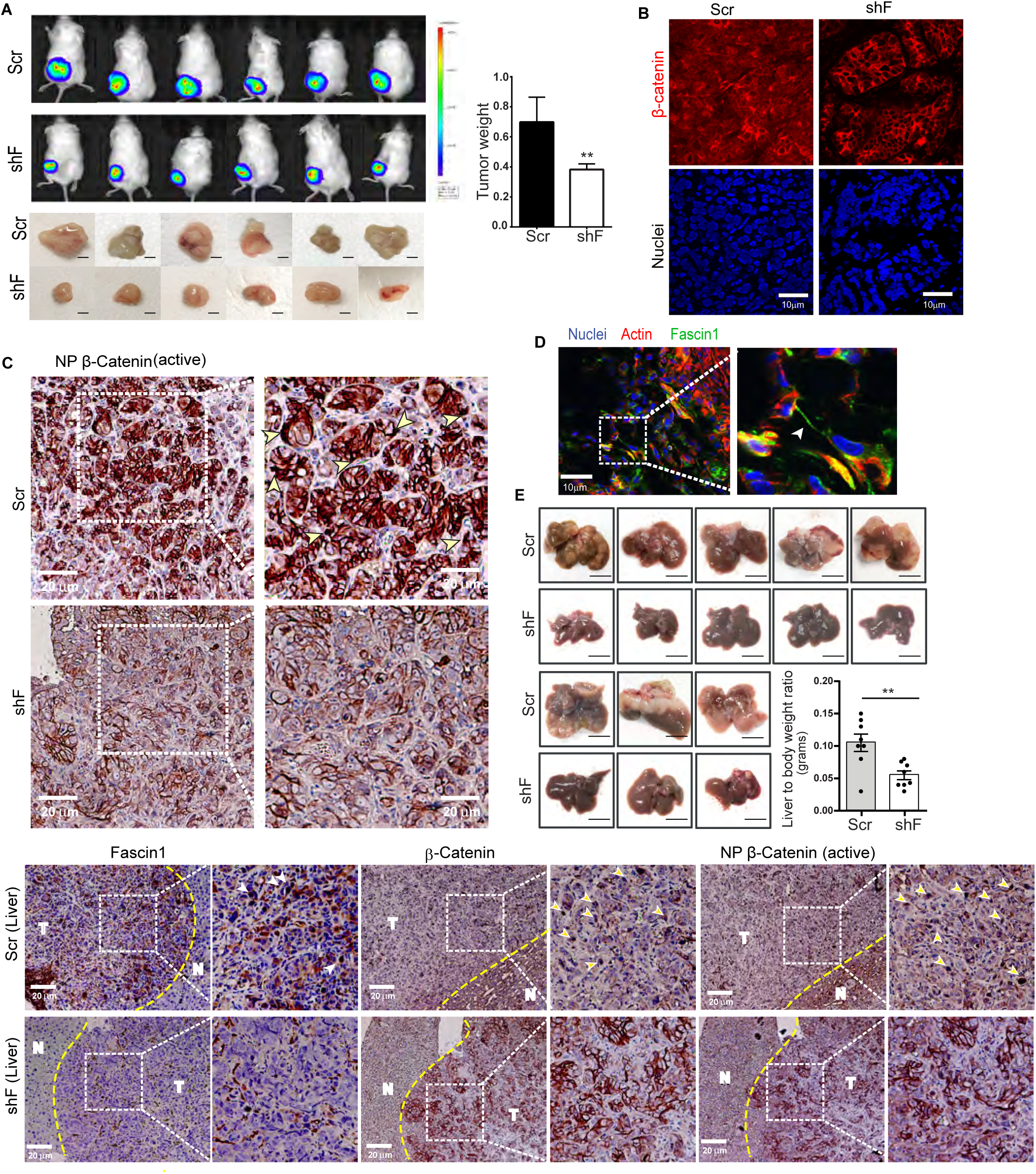
Fascin1’s function in Wnt/β-catenin signaling accelerate and amplifies primary and secondary tumor growth in mice. **(A)** IVIS imaging of NSG mice injected subcutaneously with HT-29/19A-Scr and shF cells. Fascin1 KD significantly decreased primary tumor weight. Asterisk (**) denotes p<0.001, n=6. **(B)** Confocal immunofluorescence shows distribution of total β-catenin in primary tumors of mice injected with Scr and shF cells. Nuclei were counter stained with DAPI. Scale bar, 10 μm. **(C)** Immunohistochemistry shows distribution of active (NP) β-catenin in primary tumors from mice injected with HT-29/19A-Scr or shF cells. Yellow arrowheads label nuclear accumulation of active β-catenin. **(D)** Confocal image of primary tumor from Scr injected mice shows TM like remodeled AJs and the distribution of actin (red) and fascin1 (green) in these remodeled cell-cell junctions (arrowhead). Nuclei were counter stained with DAPI. Right panel is a magnification of the boxed area. **(E)** Secondary tumor growth in liver of mice injected with Scr or shF cells. Asterisk (**) denotes p<0.0.001, n=8. Immunohistochemistry of secondary tumors from mice injected with Scr and shF cells shows distribution of fascin1, total β-catenin and active (NP) β-catenin in tumor (T) and surrounding normal (N) tissue. White arrowheads identify nuclear accumulation of β-catenin.

### In CRCs fascin1 expression is linked to hybrid EMT

Since fascin1 remodels AJ mechanotransduction, we postulated that fascin1 upregulation in CRCs could also influence signals regulating EMT. More recently, fascin1 overexpression in epithelial ovarian cancers has been linked to increased SNAI1 levels suggesting fascin1 upregulation could have a direct effect on EMT.(Li et al., 2018) For this we analyzed RNA-Seq data from CRCs profiled by the TCGA consortium, which revealed a significant inverse correlation between mRNA expression of fascin1 and the epithelial phenotype stability factors (PSFs) *OVOL1* and *GRHL2* (Fig 7A) and conversely, a significant positive correlation with the metastable markers *SNAI1/2/3*.(Jolly et al., 2016) The modest inverse correlation of fascin1 with *OVOL* and *GRHL2* suggests dampening of epithelial characteristics but not complete EMT and the positive correlation with *SNAI* suggests stabilization of the hybrid epithelial-mesenchymal (E/M) state. We also identified a significant positive correlation between fascin1 and *ZEB1/2* expression. Since stromal cells (e.g. fibroblasts) also express fascin1 and *ZEB1/2* proteins, we also performed these analyses in RNA-Seq data obtained from 61 CRC cell lines, which revealed similar findings confirming these changes were tumor intrinsic (Fig 7B). Finally, we examined the expression of these TFs in the normal mucosa and intestinal tumors derived from mutant *Apc* mice, which revealed a significant upregulation of fascin1 and *Snai3* mRNA in intestinal tumors but no change in the levels of *Ovol1*, *Grhl2* or *Zeb1/2* (Fig 7C). Together, these data are consistent with fascin1’s function in CRC cell AJ remodeling.

**Figure 7.**
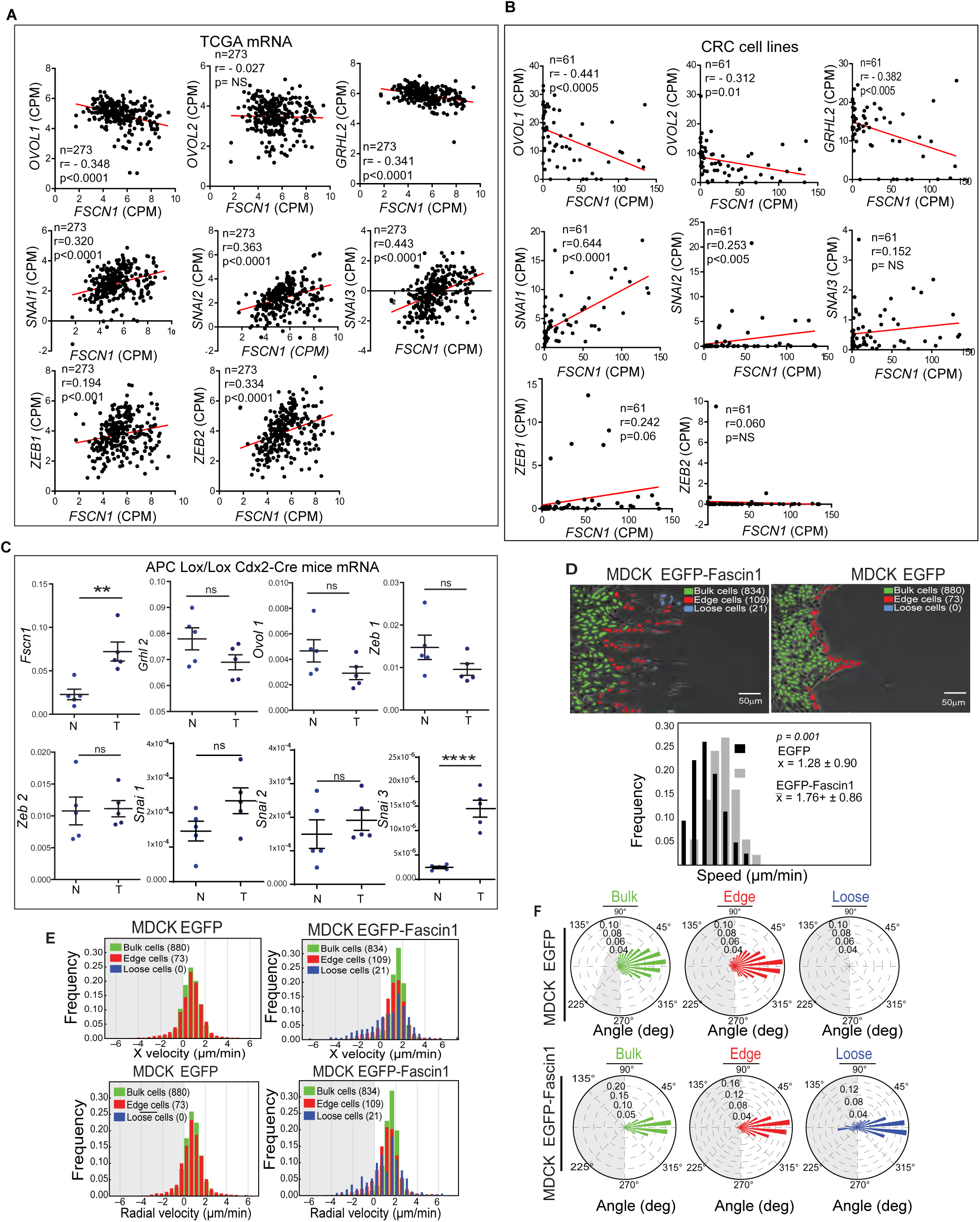
Fascin1 expression in CRCs correlates with a hybrid E/M metastable state. **(A-B)** Correlation analysis of mRNA expression levels of fascin1 with epithelial (*OVOL1*, *OVOL2* and *GRHL2*), mesenchymal (*ZEB1* and *ZEB2*), and E/M stability markers (*SNAIL1*, *SNAIL2*, *SNAIL3*) in **(A)** CRCs, n=273, and **(B)** CRC cell lines, n=61. **(C)** Analysis of mRNA expression levels of fascin1; (OVOL1 and GRHL2), mesenchymal (ZEB1 and ZEB2), and E/M stability markers (SNAIL1, SNAIL2, SNAIL3) in intestinal tumors and normal intestinal tissue from mutant Apc mice. Asterisks denote the following: **p<0.001, n=5 and ****p<0.0001, n=5. **(D)** Phase contrast image of scratch wound of MDCK control cells expressing EGFP or overexpressing EGFP-Fascin1. Cells identified by our tracking method are shown in green (bulk), red (edge) or blue (loose). Scale bar, 50 μm. See methods for description of group classifications. Quantification of instantaneous migration speeds are shown as mean 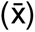 ± SD. P-value from Mann-Whitney U test comparing MDCK EGFP and MDCK EGFP-Fascin1 cells. **(E)** Quantification of instantaneous migration velocity in X coordinates only and in XY-radial coordinates. Instantaneous migration velocity frequencies in XY-radial coordinates. **(F)** Instantaneous velocity-angle histograms of migrating MDCK EGFP and MDCK EGFP-Fascin1 bulk, edge, and loose cell clusters.

### Fascin1 regulates “collective plasticity” and bidirectional cell migration

As collective cell migration is a hallmark of the hybrid E/M metastable state we assessed the role of fascin1 in a wound-closure assay. To determine how AJ remodeling by fascin1 could modulate the behavior of migrating cells, we used a reductionist approach and elected to overexpress fascin1 in the non-transformed epithelial cells, MDCK. Intestinal epithelial cells move collectively as closely adherent sheet of migrating cells. As expected, EGFP expressing MDCK cells migrated as a sheet of tightly adherent cells very similar to normal intestinal epithelial cells (Supplementary Videos 3-4; Fig 7D). As predicted, fascin1 overexpression significantly increased cell migration rates compared to control cells. However, unlike control cells, fascin1 expressing cells migrated as loosely adherent groups of cells resembling cell migration patterns previously described as “collective plasticity.”(Friedl and Mayor, 2017) We identified three groups of migrating EGFP-Fascin1 cells that we refer to as loose, edge and bulk, which migrated at different speeds and with different patterns of migration (Supplementary Video 3-4; Supplementary Table 1; Figs 7D-7F). Loose and edge cells migrated using TM-like structures of variable lengths connecting the back of a leading cell with the front of a follower cell (Supplementary Videos 3-4, Supplementary Fig 6A). By measuring the area of migrating bulk cells, it is clear that migrating bulk cells also remodel cell-cell adhesions although the TM-like cell-cell junctions are much smaller (Supplementary Videos 3 and 4; Supplementary Fig 6B). Migrating EGFP-Fascin1 cells displayed large gaps within the migrating cell clusters, indicative of increased tension at cell-cell adhesion sites and continuous AJ remodeling during collective cell migration (Supplementary Videos 3 and 4; Supplementary Fig 6D). Together these data demonstrate that fascin1’s AJ remodeling property also influences the behavior of collectively migrating epithelial cells. Additionally, we made the most striking finding namely, that fascin1 overexpressing loose cells displayed significantly more backwards migration events in the direction opposite from the wound edge, which were quantitatively measured by the shape metrics (Supplementary Video 4; Figs 7D-7F, Supplementary Fig 6C, Supplementary Table 2). Based on the analysis of X and XY (radial) coordinates, the backwards migration of EGFP-Fascin1 cells was significantly different from the consistent forward migration of control EGFP cells. To the best of our knowledge fascin1’s role in bidirectional migration has not been reported before. And to the best of our knowledge this is also the first identification of actin-binding protein that regulates bidirectional cell migration. Villin1 is an actin bundling protein that assembles filopodia and increases intestinal cell migration rates during normal intestinal wound repair.(George et al., 2013; Ubelmann et al., 2013) MDCK cells overexpressing EGFP-Villin1 migrated unidirectionally as a sheet of tightly adherent cells resembling normal intestinal epithelial cell migration, demonstrating the unique ability of fascin1 to produce bidirectional “collective plasticity” (Supplementary Video 5).

## Discussion

High fascin1 expression in CRCs is an independent negative prognostic factor for survival and increases the risk of disease recurrence and death by seven-fold.(Tampakis *et al*., 2021) A role for fascin1 as a driver of colon tumorigenesis was shown previously by overexpression of fascin1 in the CRC cell line HT-29 which increased metastatic lesions in mice by ten-fold. (Vignjevic *et al*., 2007) Similarly, constitutive overexpression of fascin1 in the intestinal epithelium in the *Apc*-mutated mouse background has been shown to accelerate tumor initiation and progression in this mouse model of colon cancer. (Schoumacher et al., 2014) It is clear from these human and animal studies that fascin1 is both a driver and prognostic marker of aggressive clinical features of CRCs. Our study demonstrates statistically significant correlation between high fascin1 expression and high-grade colon cancers. In this study we provide for the first time, a molecular mechanism for fascin1’s function in colorectal tumor growth and metastasis. Mechanical cues can initiate tumor-like gene expression pattern in pre-neoplastic tissues of the *Apc* mutant mice, this could explain why conditional expression of fascin1 in mutant Apc mice promotes tumor initiation.(Whitehead *et al*., 2008) This study also lends support to our hypothesis that fascin1’s function as a mechanosensor regulates colorectal carcinogenesis. In the absence of Wnt ligands, mechanical forces that change AJ mechanotransduction are sufficient to promote nuclear accumulation of β-catenin, and in this fashion induce mechanical-strain induced changes in epithelial cell behavior.(Benham-Pyle et al., 2015) Mechanosensitive activation of Wnt/β-catenin signaling does not require oncogenic transformation. For instance, in the quiescent epithelium, mechanical strain is sufficient to activate β-catenin and cell cycle re-entry.(Benham-Pyle *et al*., 2015; Fernandez-Sanchez *et al*., 2015; Roper et al., 2018) We propose that the profound changes in AJ actin cytoskeleton and AJ mechanotransduction that are induced in response to fascin1 expression in epithelial cells are sufficient for mechanosensitive Wnt/β-catenin signaling and to generate oncogenic characteristics. Using the TOPFLASH reporter plasmid it has previously been reported that the fascin1 gene promoter is activated by the β-catenin/TCF signaling complex. (Vignjevic *et al*., 2007) While we did not see a similar effect of Wnt signaling on fascin1 expression, a feedback mechanism between Wnt and fascin1 is possible. The nuclear pore complex is also a mechanosensor that is regulated by nuclear envelope (NE) tension which modulates the nuclear pore diameter to regulate the nuclear trafficking of transcription factors.(Dahl et al., 2008; Elosegui-Artola et al., 2017; Fichtman and Harel, 2014; Wolf, 2009) In a breast cancer cell line the NE protein nesprin-2 giant has been shown to interact directly with fascin-1 and this association was shown to regulate nuclear deformation during cell motility.(Jayo et al., 2016) Their study also demonstrated that this function of fascin1 was independent of its actin binding property.(Jayo *et al*., 2016) We hypothesize that if this is also true in CRC cells, then one pool of fascin1 could bundle actin and remodel AJ mechanotransduction, while the second pool of fascin1 could modulate NE tension to facilitate nuclear trafficking of β-catenin. The role of mechanical cues on tumor growth and progression is well documented and we propose that fascin1 functions as a mechanosensor to drive colorectal carcinogenesis.(Dombroski et al., 2021; Huang et al., 2019; Simi, 2015)

In this study we show that fascin1 expression in CRC cells profoundly transforms AJ-associated actin cytoskeleton rearranging it perpendicular to the cell-cell junctions (Fig 8).This also results in the rearrangement of actomyosin contractility which now pulls on the AJ-associated actin bundles from both sides of the cell-cell junctions, generating discontinuous, dynamic adherens junctions (Fig 8). Our study shows that in this manner by simply sorting AJ-associated mechanosensors (i.e. α-actinin, EPLIN, α-catenin, vinculin) fascin1 can generate substantial changes in AJ mechanotransduction. It is known that fascin1 and α-actinin form actin bundles with very different inter-filament spacing, 8 nm *versus* 35 nm, respectively.(Winkelman *et al*., 2016) We propose that in CRC cells, the tightly packed actin bundles formed by fascin1 exclude non-muscle myosin II (NMII) which is revealed by the absence of fascin1 from the contractile actin bundles of the AJs (Fig. 4B). We suggest that this is the likely mechanism that regulates the sorting of α-actinin, EPLIN and fascin1 cross-linked actin networks at the AJs of colon cancer cells. It is also known that both fascin1 and α-actinin assemble stable cross-linked actin networks but when these two structures are mixed together they form unstable structures because α-actinin crosslinks impose unfavorable and energetically costly defects in hexagonally packed structures formed by fascin1 and *vice versa*.(Freedman et al., 2019) We propose that such energetically costly defects caused by fascin1 expression in CRC cells is responsible for increased AJ actin dynamics (Fig 4E) and increased AJ plasticity during collective cell migration (Supplementary Fig 7). Targeted Wnt therapies for the treatment of CRCs have been unsuccessful due to the pleiotropic effects of Wnt. Most of these drugs also target ligand-receptor Wnt signaling, a likely factor in their ineffectiveness. The fascin1 knockout mice do not have any major defects, and since fascin1 is absent from the normal colon, our study also identifies fascin1 for targeted therapies for CRCs.

**Figure 8.**
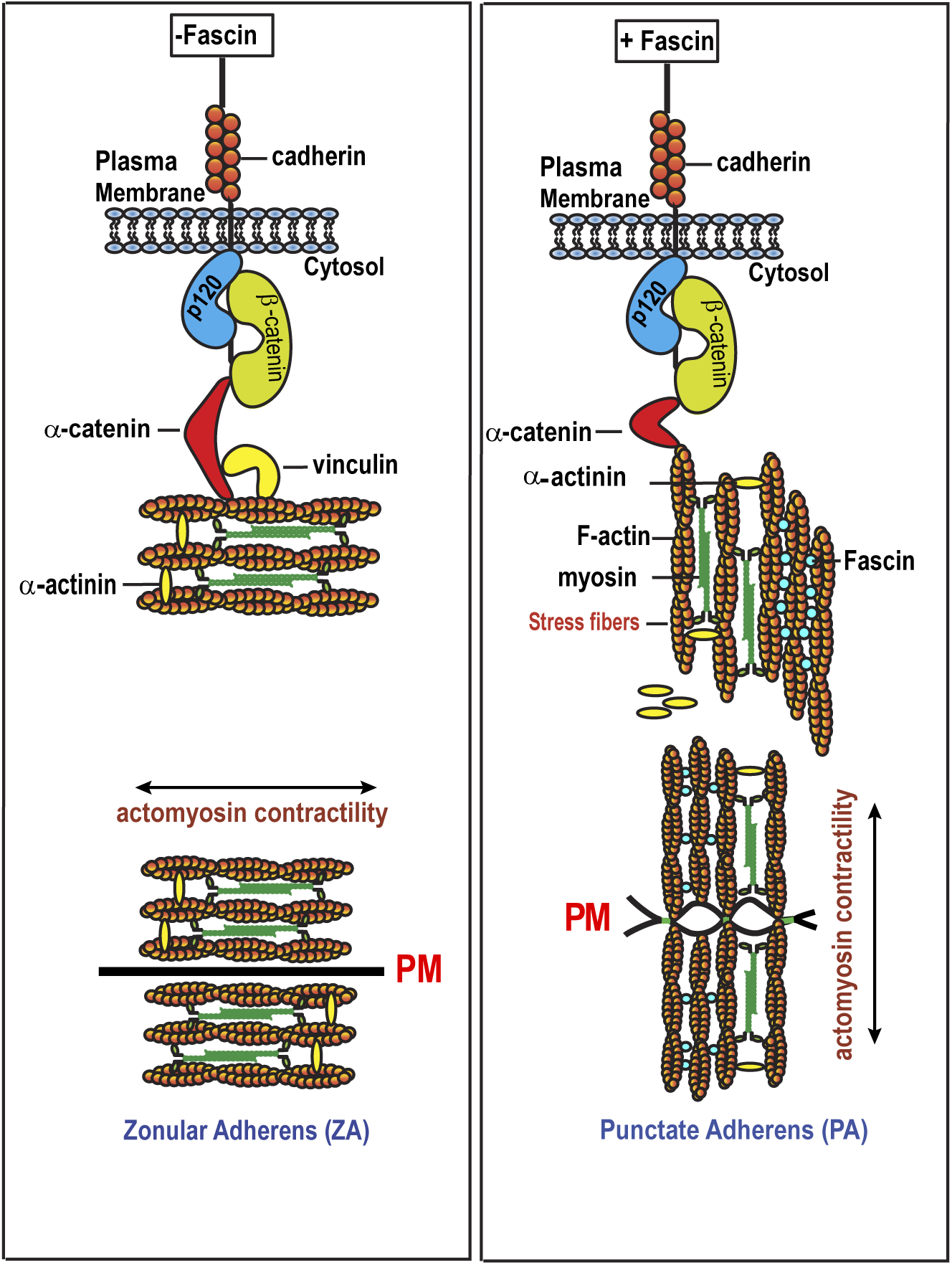
Fascin remodels CRC cell AJs. In normal epithelial cells, AJ-associated actin cytoskeleton is aligned parallel to the plasma membrane (PM). fascin1 expression in CRC cells profoundly transforms AJ-associated actin cytoskeleton rearranging it perpendicular to the cell-cell junctions. Actomyosin contraction of RABs pulls on opposite sides of the PM making PAs discontinuous with gaps in cell-cell junctions. Actomyosin contraction of CABs pulls in direction of PM sealing cell-cell junctions, making them continuous and stable.

In our study, we also identify the ability of fascin1 to assemble TM-like ultra-long, large, and long-lived cell-cell adhesions. We show that these TM-like structures contain E-cadherin and β-catenin, which suggests that the origins of these structures are similar to epithelial protrusions that assemble and/or repair AJs in normal intestinal epithelial cells.(Li et al., 2020) Our findings indicate that fascin1 appropriates this molecular machinery to construct these TM-like remodeled AJs. We show that in CRC cells these super-elongated cell-cell junctions share many structural and functional similarities with TNTs, neuronal growth cones and TMs such as their morphology, their inclusion of tubulin and organelles, and their ability to hover over cell surfaces while connecting distant cells.(Osswald *et al*., 2015; Rustom *et al*., 2004) TNTs are understood to correspond to invadopodia *in vivo* and TMs *in vivo* have been shown to regulate tumor cell migration/invasion but also tumor proliferation and radioresistance.(Naphade et al., 2015; Osswald *et al*., 2015) While these TM-like structures remain to be identified in CRCs, we hypothesize that this property of fascin1 also implicates it in CRC cell invasion and chemoresistance. These TM-like intercellular connections could have other functions e.g. as conduits for cell signaling allowing cells at the invasive front and the tumor center to communicate directly, or to function like cytonemes to traffic morphogens and generate morphogen gradients. Since stromal cells also expression fascin1, morphogen trafficking between the stromal and tumor cells can be envisioned and could influence CRC tumor heterogeneity. Tumor heterogeneity is responsible for the failure of most cancer therapies and this may be the molecular basis for fascin1’s strong association with treatment-refractory CRCs.(Ristic *et al*., 2021)

The expression of TFs such as *OVOL* and *ZEB* and *OVOL* and *SNAI* induce the hybrid E/M metastable state.(Jia et al., 2017) We find that fascin1 expression in CRCs is linked to the hybrid E/M state which is consistent with its role in AJ remodeling. While we predict that the majority of carcinomas expressing fascin1 will have an E/M phenotype, some fascin1 expressing cancers could reflect a complete phenotypic transition to M due to rewiring of EMT regulatory networks and/or differences in epigenetic reprogramming. Our cell migration studies demonstrate that fascin1 expressing cells migrate collectively using TM-like remodeled cell-cell connections which likely function *in vivo* as guidance cues. This is supported by the knowledge that similar AJ elongation during morphogenesis provides directional cues to the migrating cells.(Niewiadomska et al., 1999) Neural crest cells in developing embryos, astrocytes, and glioma cells also form filamentous TM or TM-like cell-cell interactions while migrating as loosely cohesive cell clusters.(Osswald *et al*., 2015; Scarpa *et al*., 2015) It is noteworthy that fascin1 is expressed in all these cell types.(Gunal et al., 2008; Zhang *et al*., 2008) This pattern of “collective plasticity” and the use of TM-like remodeled AJs during cell migration is conserved during morphogenesis and tumorigenesis. Based on that we interpret our findings as reactivation of embryonic cell migration patterns in CRCs, and we suggest that this may be a recapitulation of fascin1’s function in the developing gut. Such patterns of cell migration are not seen during normal intestinal wound healing and in our study they were not duplicated by the intestinal actin bundling protein villin1 which may explain why fascin1 is re-expressed during colorectal carcinogenesis. We hypothesize that these TM-like structures could provide tumor cells with an advantage when responding to external cues by rapidly remodeling AJs allowing CRC cells to switch between more tightly or more loosely adherent cells as they navigate their way through the underlying tissue and during extravasation and intravasation.(George *et al*., 2013) Normal epithelial cell migrate unidirectionally. However, human mesenchymal stem cells migrate bidirectionally in response to chemokine signaling.(Smith et al., 2012) Similarly, bidirectional migration of leukocyte T cells and vertebrate axon growth cones is regulated by chemoattraction and chemorepulsion.(Poznansky et al., 2000; Vianello et al., 2005) We hypothesize that fascin1’s function in bidirectional migration could enhance the response of tumor cells to chemoattractants and chemorepellents thereby augmenting the metastatic potential of CRC cells. Bidirectional migration of CRC cells could also prevent spatial competition. In summary, this study provides for the first time a molecular mechanism for fascin1 in colorectal carcinogenesis and identifies fascin1 as a mechanosensor that remodels AJs to drive colorectal tumor growth and metastasis.

## Grant Support

This study was supported by National Institutes of Diabetes and Digestive Kidney Diseases (grant DK-117476 to S.K.); and the Public Health Service (grant DK-56338).

## Abbreviations

AJ: adherens junctions
CRC: colorectal carcinoma
E/M: hybrid epithelial-mesenchymal
GI: gastrointestinal
nAJ: nascent adherens junctions
SSL: sessile serrated lesions
TJ: tight junction
TM: tumor microtube
TNT: tunneling nanotube

## Author Contributions

Eric Pham, Amin Esmaeilniakooshkghazi, Sudeep P. George, Afzal Ahrorov, Fabian R. Villagomez, Michael Byington, Srijita Mukhopadhyay, Srinivas Patnaik, Monali Naik and Saathvika Ravi, Niall Tebuttt, Jennifer Mooi, Camilla M. Reehorst performed the experiments described in this study. Jacinta C. Conrad analyzed the migration data and provided important guidance for the experimental design of these studies. John M. Mariadason is a long-standing collaborator of the contributing author and provided the patient cohort data, analyzed the patient cohort data as well as provided valuable suggestions during the writing of this manuscript.

## Data Transparency

Data, analytical methods, and study material will be made available to other researchers once the study has been completed and published.

## Figure Legends

**Supplementary Figure 1. (A)** Endogenous fascin1 expression in HT-29/19A-Scr and HT-29/19A-shF cells and ectopic overexpression of EGFP-Fascin1 in HT-29/19A-shF cells. Tubulin was used as a loading control. This immunoblot is a representative of at least three independent experiments with similar findings. **(B)** Confocal immunofluorescence shows distribution of F-actin (red), tubulin (green) and fascin1 (magenta) in nAJs of Scr cells. Nuclei were counter stained with DAPI. Scale bar, 30 μm. **(C)** Phase contrast image of HT-29/19A-Scr and shF cells. Scale bar, 10 μm. Rate of elongated nAJ assembly was measured in HT-29/19A-Scr and shF cells. Data are presented as mean ± S.E.M.; Student’s t-test. Asterisk (***) denotes p<0.001, n=10 fields/group and >300 cells/group. **(D)** Phase contrast image of HT-29/19A-Scr cells. Scale bar, 10 μm. Rate of elongated nAJ assembly was measured in Scr cells 48 h post-treatment with vehicle control (DMSO), cytochalasin B (250 nM), or colchicine (1 mM). Data are presented as mean ± S.E.M.; Students’s t-test. Asterisk (***) denotes p<0.001, n=10 fields/group and >300 cells/group. Asterisk (****) denotes *p<0.0001,* n=10 fields/group and >300 cells/group. **(E)** Ectopic overexpression of EGFP-Fascin1 in MDCK cells. Tubulin was used as a loading control.

**Supplementary Figure 2. (A)** Immunoblot shows ectopic overexpression of EGFP and EGFP-fascin1 in HT-29 cells, a CRC cell line that lacks endogenous fascin1 protein. Tubulin was used as a loading control. This immunoblot is a representative of at least three independent experiments with similar findings. **(B)** Confocal immunofluorescence of HT-29 EGFP and HT-29 EGFP-Fascin1 cells shows distribution of AJ associated actin (red) and EGFP (green). Nuclei were counter stained with DAPI. Scale bar, 10 μm. **(C)** Confocal image of MDCK EGFP and MDCK EGFP-Fascin1 cells shows distribution of AJ associated actin (red) and EGFP (green). Nuclei were counter stained with DAPI. Scale bar, 10 μm. **(D)** Confocal image of shF cells overexpressing EGFP or EGFP-Fascin1 shows distribution of AJ associated actin (red) and EGFP (green). Scale bar, 10 μm. **(E)** Confocal image shows the distribution of focal adhesion associated actin (red), α-actinin (green) and vinculin (green) in Scr and shF cells. Right panels are magnifications of the boxed areas. Immunoblot shows total vinculin, fascin1 and α−actinin levels in Scr and shF cells. Tubulin was used as a loading control. This immunoblot is a representative of at least three independent experiments with similar findings.

**Supplementary Figure 3. (A)** Immunoblot shows total E-cadherin and β-catenin levels in Scr and shF cells. Tubulin was used as a loading control. This immunoblot is a representative of at least three independent experiments with similar findings. **(B)** Confocal immunofluorescence z-stack of MDCK EGFP-Fascin1 cells shows distribution of fascin1 and β-catenin. Nuclei were counter stained with DAPI. Scale bar, 20 μm. Data obtained from n = 10 fields/group, and >15 cells/group. (**C)** ChIP-qPCR with β-catenin antibody and cell lysates from HT-29/19A-Scr and shF cells. ChIP with IgG and negative control primers against a region of MMP9 were used as control. Data are representative of n=3 biological replicates, mean ± S.E.M.; Student’s t-test. Asterisk (*) denotes *p<0.05, n=3. NS, not significant, **(D)** Wnt signaling activity in *BRAF^V600E^* mutant HT-29/19A-Scr and shF cells was monitored using TOP Flash luciferase reporter assay. Data are shown as mean ± S.E.M., Student’s t-test. Asterisk (*) denotes p<0.01, n=3.

**Supplementary Figure 4. (A)** Confocal image of shF EGFP and shF EGFP-Fascin1 cells shows xz distribution of actin (red) and EGFP (green). Nuclei were counter stained with DAPI. Scale bar, 10 μm. **(B)** Confocal image of MDCK EGFP and MDCK EGFP-Fascin1 cells shows xz distribution of actin (red) and EGFP (green). Nuclei were counter stained with DAPI. Scale bar, 10 μm. **(C)** Confocal image shows xz distribution of actin (red) in HCT-116 expressing scrambled shRNAs (Scr) and HCT-116 cells transduced with shRNAs to KD fascin1 (shF cells). Nuclei were counter stained with DAPI. Scale bar, 10 μm. Immunoblot shows total fascin1 levels in HCT-116 cells expressing scrambled shRNA (Scr) and HCT-116 with fascin1 KD using fascin1 specific shRNA (shF). Actin was used as a loading control. This immunoblot is a representative of at least three independent experiments with similar findings.

**Supplementary figure 5. (A)** Confocal immunofluorescence of primary tumors from mice injected with Scr or shF cells labeled for cell proliferation (Ki67, green) and apoptosis (cleaved caspase-3, red). Nuclei were counter stained with DAPI. Scale bar, 10 μm. **(B)** Scr and shF cells labeled with dye (red) injected into 2-day old zebrafish embryos and metastasis scored for 72 h post-injection. This is a representative of three independent experiments with similar findings.

**Supplementary figure 6 (A)** Phase contrast microscopy shows TM-like long intercellular junction connecting two migrating MDCK EGFP-Fascin1 loose cells. Arrowheads indicates the TM-like structure. **(B)** Total area (μm^2^) covered by ‘bulk’ cells was measured in migrating MDCK EGFP and MDCK EGFP-Fascin1 cells. Asterisk (****) denotes p<0.0001, n=3. **(C)** Time lapse phase contrast imaging of MDCK EGFP-Fascin1cells captured every 10 min. Start time, 20.30 and finish time, 1:02:40 **(D)** Large gaps (arrowheads) identified in migrating MDCK EGFP-Fascin1 cells.

## Methods

### Cells and plasmids

HT-29/19A, HCT-116 and MDCK cells are described by us previously.(Mathew et al., 2008; Patnaik et al., 2016) MRC-5, HT-29 cells were purchased from American Type Culture Collection (ATCC, Rockville, MD). Human fascin1 plasmid was a kind gift from Dr. Shigeko Yamashiro (Rutgers University, New Brunswick, NJ). HT-29/19A and HCT-116 cells were transduced with lentivirus expressing short-hairpin RNAs (shRNAs) targeting fascin1 (Sigma-Aldrich, St. Louis, MO). For the rescue experiments, EGFP-Fascin1 plasmid without 3’UTR region were overexpressed in HT-29/19A-shF cells. EGFP-Fascin1 or EGFP were also overexpressed in HT-29 cells that lack endogenous fascin1 protein.

### Tumor spheroid culture

Tumor spheroids were generated as described previously.(Humtsoe et al., 2012) Briefly, cells were grown on standard tissue culture dishes to 80% confluence, and re-plated 24 h prior to initiating tumor spheroids. For tumor spheroids 1 x10^6^ cells/mL were plated on agarose-coated dishes. Spheroids were imaged using a Nikon Eclipse TE2000-U inverted microscope equipped with a CoolSnap ES charge-coupled device (CCD) camera. For hanging drop cultures, 2000 cells in 20 μl drops were plated on the inside lids of 10 cm dishes, and the lids were inverted prior to incubation.

### Immunoblotting

Cell lysates were prepared as described previously.(Wang et al., 2016) Briefly, proteins were separated by SDS-PAGE, transferred to nitrocellulose membranes, immunoblotted and detected with ECL. All Western blots are representative of at least three independent experiments with similar findings.

### Cell migration assay

Migration assay was performed as described previously.(George *et al*., 2013) Briefly, cells were plated on 6-well tissue culture treated dishes and allowed to grow to confluence. Upon confluence, the monolayers were wounded with a sterile blade. Wells were rinsed with phosphate buffered solution (PBS), and cells were treated with MRC-5 conditioned media (diluted in 3 volumes 10% FBS DMEM). Cells were treated for 20 h prior to imaging.

### Scratch-wound image analysis

For image analysis, cells were identified through a series of filters applied to individual images using Python, which produced binary images. Adjacent frames were shifted to maximize the total overlap in the binary images and cells were assigned to tracks based on maximum overlap. Cells unaccounted for by this method were assigned tracks by minimizing the centroid displacement between remaining cells. Edge cells were identified as being within 75 μm of the migratory front. Loose and bulk cells were identified by their position relative to the migratory front and edge cells. Instantaneous centroid velocities were used for further analysis. Cell tracks that were detected in a minimum of 10 frames were analyzed.

### Cell-cell junction remodeling assay

For mature AJ remodeling assays, 5×10^4^ cells were plated in 6-well tissue culture treated dishes and allowed to form small colonies (<40 cells). The cells were treated with MRC-5 conditioned media, and imaged every 5 minutes. Images were taken with a Nikon Eclipse TE2000-U inverted microscope and CoolSnap ES CCD camera.

### Invasion Assay

For invasion assays, tumor spheroids formed by the hanging drop method were isolated 96 hours after suspension culture, and embedded in a Collagen I matrix (2mg/mL). The Collagen I matrix was allowed to solidify for 90 minutes prior to being overlaid with 10% FBS containing DMEM with 50 ng/mL HGF. Additionally, the tumor spheroids were treated with vehicle control (DMSO). The invasion occurred over 8 days.

### Soft agar colony formation assay

First a layer of 0.5% agarose-containing DMEM was solidified at the bottom of wells of 6-well dishes to prevent cells from attaching. Cells were mixed into liquid 0.33% agarose at the desired cell numbers and allowed to solidify on top of the 0.5% layer. For HT-29/19A cells 500 cells were used. Colonies greater than 0.05μm^2^ in area were counted.

### Mice

NOD-*scid* IL2Rgamma^null^ mice (NSG) (Jackson Laboratory, Bar Harbor, ME) were injected subcutaneously with 2 x 10^6^ Scr and shF cells and intra-splenic with 1×10^6^ Scr and shF cells. Three weeks’ post-injection primary and secondary tumors were excised and measured.

### Zebrafish metastasis assay

Zebrafish (Danio rerio) were reared and maintained at 28.5°C. *Tg(kdrl:EGFP)mitfa^b692/b692^* was generated by crossing *Tg(kdrl:EGFP)* with *mitfa^b692/b692^* (Zebrafish International Resource Center, Eugene, OR). 300-500 HT-29/19A Scr and HT-29/19A-shF cells labeled with CM-Dil (ThermoFisher) were microinjected into yolk sac of 2-day old embryos using a pressure injector (Harvard Apparatus, Holliston, MA) and manipulator (MM33-Righ, Märzhäuser Wetzlar, Germany). Embryos and cells injected in the yolk were incubated at 34°C and imaged using OLYMPUS IX51 fluorescence microscope, OLYMPUS XM10 camera and cellSens Dimension software (Olympus, Center Valley, PA, USA).

### Immunofluorescence and live cell fluorescence microscopy

Confocal images were acquired using the Olympus FV2000 inverted confocal microscope. Unless otherwise indicated all data were obtained by performing three independent experiments with n = 10 fields/group and >300 cells/group were analyzed each time. Nuclei were counter stained with DAPI. Tumor spheroids were studied in three independent experiments with 10 spheroids per group in each experiment. Plot profiles were generated by measuring the distribution of AJ associated proteins in regions of interest (ROI) highlighted by a white line, 10 μm long.

### Chromatin Immunoprecipitation

ChIP was carried out as previously described.(Bado et al., 2016) Briefly, cells were fixed with 1.5% paraformaldehyde, lysed, nuclei were purified, and then chromatin was sheared by sonification. Immunoprecipitation was carried out with β-catenin monoclonal antibody and Protein G magnetic beads (Sigma-Aldrich). For PCR, the following primers were used:

MMP9:
F: 5’-CCTCCTTAAAGCCCCCACAA-3’,
R: 5’-CAGTCCACCCTTGTGCTCTT-3’;
CCND1:
F: 5’-CAGAAGAGCGCGAGGGAG-3’,
R: 5’-ATGGAACACCAGCTCCTGTG;
MYC,
F: 5’-GAGGCGAACACACAACGTCTT-3’,
R: 5’-CGCAACAAGTCCTCTTCAGAAA-3’
MMP9 negative control,
F: 5’-GTGGGACCTCAACGTCTGTC-3’,
R: 5’-CACCACCTCTAAGCACTGACAT-3’.
Fascin1:
F: 5’-AGGCGGCCAACGAGAGGAAC-3’
R: 5’-ACGATGATGGGGCGGTTGAT’3’
CD44:
F: 5’-GCAGTCAACAGTCGAAGAAGG’3’
R: 5’-TGTCCTCCACAGCTCCATT-3’
MMP7:
F: 5’-CTTACCTCGGATCGTAGTGG-3’
R: 5’-CCCCAACTAACCCTCTTGAAGT-3’
AXIN2:
F: 5’-AACCTATGCCCGTTTCCTCTA-3’
R: 5’-GAGTGTAAAGACTTGGTCCACC-3’
TCF1:
F: 5’-CCAGTGTGCACCCTTCCTAT-3’
R: 5’-AGCCCCACAGAGAAACTGAA-3’

### Promoter reporter assays

**Promoter reporter assays.** Cells were transfected in 24-well dishes using Lipofectamine 2000 (ThermoFisher) with β-catenin reporter Super 8X TOPFlash and pSV-β-Galactosidase control vector. All cells were lysed approximately 24 hours post-transfection, and luciferase activity was measured with Luciferase Assay (Promega, Madison, Wisconsin, USA) and β-Galactosidase with chlorophenol red-β-D-galactopyranoside (CPRG). Reporter activity was normalized to β-Galactosidase activity. 24 hours’ post-transfection, cells were treated with DMSO, 20µM G2, or 100µM G2 for 24 hours. Luminescence was quantified with VICTOR Multilabel Plate Reader (PerkinElmer, Waltham, MA, USA).

### Fascin1 and β-catenin immunohistochemistry scoring

Patient tumor microarrays were immunostained for fascin1 and total β-catenin. Fascin1 was scored using the method of Hashimoto *et al* and β-catenin was scored by adapting the method of Hugh *et al*.(Hashimoto *et al*., 2006; Hugh et al., 1999) Immunohistology in paired tumor panels show expression of fascin1 and β-catenin in the same tumor.

### Statistical Analysis

Data are expressed as mean ± standard deviation (SD) or ± standard error (SE). Significance was determined by Student’s t-test, Mann-Whitney U test, Kuiper’s test, or Watson U2 test.

### Study approval

All animal studies were performed in accordance with animal protocols approved by the University of Houston Institutional Animal Care and Use Committee.

***All authors had access to the study data and have reviewed and approved the final manuscript.***

## Notes

### Competing Interest Statement

The authors have declared no competing interest.

### Summary of Updates

In this manuscript, we show how fascin1 expression in colon cancer cells results in the competitive exclusion of AJ associated actin-binding and actin-bundling proteins resulting in remodeling of AJ associated actomyosin contractility and activation of force-sensitive oncogenic signaling. During morphogenesis and during cancer cell invasion a flexible type of collective cell migration referred to as "collective plasticity" occurs where some properties of single cell migration is acquired by collectively migrating cells, when sheets of cells and detached groups of cells maintain cell-cell adhesions while migrating as a loosely cohesive group. The mechanisms regulating "collective plasticity" remains unclear. Moreover, the integration of guidance cues between such functionally and morphologically distinct cell populations are yet-to-be-defined. Moreover, the proteins regulating the remodeling and weakening of AJs during "collective plasticity" remain unidentified. In this manuscript, we show how fascin1's AJ remodeling function induces "collective plasticity". We demonstrate the unique function of fascin1 is assembling tumor microtubule-like cell-cell connections which provide guidance cues during "collective plasticity". We show for the first time, that unlike other actin-bundling proteins associated with collective cell migration, fascin1 is unique in generating bidirectional cell migration. This is, to the best of our knowledge, the first identification of an actin-binding protein that regulates cell-intrinsic reverse and bidirectional cell migration. We hypothesize that fascin1's function in bidirectional migration could enhance the response of tumor cells to chemoattractants and chemorepellents thereby augmenting the metastatic potential of colon cancer cells. Bidirectional migration of colorectal cancer cells could also prevent spatial competition thus promoting tumor progression. Fascin1 is widely expressed during morphogenesis and in all types of carcinomas where its expression is always correlated with a clinically aggressive disease associated with high mortality. It is generally assumed that fascin1 regulates tumor cell migration by assembling filopodia and/or invadopodia. However, there is emerging evidence that fascin1 also has cell migration-independent functions such as the regulation of cell proliferation, oncogenesis, metastatic colonization, anoikis resistance, chemoresistance and cancer stemness although the molecular mechanism underlying these non-canonical functions of fascin1 remains unclear. Our study identifies fascin1 as a "mechanosensor" that transforms cell-cell adhesions to have a profound effect on mechanosensitive oncogenic signaling. We propose that our model where fascin1 is identified as a mechanosensor also provides a molecular mechanism for some of these non-canonical functions of fascin1.

## References Cited

Aceto, N., Bardia, A., Miyamoto, D.T., Donaldson, M.C., Wittner, B.S., Spencer, J.A., Yu, M., Pely, A., Engstrom, A., Zhu, H., et al. (2014). Circulating tumor cell clusters are oligoclonal precursors of breast cancer metastasis. Cell 158, 1110–1122. 10.1016/j.cell.2014.07.013.

Bado, I., Nikolos, F., Rajapaksa, G., Gustafsson, J.A., and Thomas, C. (2016). ERbeta decreases the invasiveness of triple-negative breast cancer cells by regulating mutant p53 oncogenic function. Oncotarget 7, 13599–13611. 10.18632/oncotarget.7300.

Barnawi, R., Al-Khaldi, S., Majed Sleiman, G., Sarkar, A., Al-Dhfyan, A., Al-Mohanna, F., Ghebeh, H., and Al-Alwan, M. (2016). Fascin Is Critical for the Maintenance of Breast Cancer Stem Cell Pool Predominantly via the Activation of the Notch Self-Renewal Pathway. Stem Cells 34, 2799–2813. 10.1002/stem.2473.

Baum, B., and Georgiou, M. (2011). Dynamics of adherens junctions in epithelial establishment, maintenance, and remodeling. J Cell Biol 192, 907–917. 10.1083/jcb.201009141.

Benham-Pyle, B.W., Pruitt, B.L., and Nelson, W.J. (2015). Cell adhesion. Mechanical strain induces E-cadherin-dependent Yap1 and beta-catenin activation to drive cell cycle entry. Science 348, 1024–1027. 10.1126/science.aaa4559.

Bidard, F.C., Peeters, D.J., Fehm, T., Nole, F., Gisbert-Criado, R., Mavroudis, D., Grisanti, S., Generali, D., Garcia-Saenz, J.A., Stebbing, J., et al. (2014). Clinical validity of circulating tumour cells in patients with metastatic breast cancer: a pooled analysis of individual patient data. Lancet Oncol 15, 406–414. 10.1016/S1470-2045(14)70069-5.

Brunet, T., Bouclet, A., Ahmadi, P., Mitrossilis, D., Driquez, B., Brunet, A.C., Henry, L., Serman, F., Bealle, G., Menager, C., et al. (2013). Evolutionary conservation of early mesoderm specification by mechanotransduction in Bilateria. Nat Commun 4, 2821. 10.1038/ncomms3821.

Cheung, K.J., and Ewald, A.J. (2016). A collective route to metastasis: Seeding by tumor cell clusters. Science 352, 167–169. 10.1126/science.aaf6546.

Cristofanilli, M., Budd, G.T., Ellis, M.J., Stopeck, A., Matera, J., Miller, M.C., Reuben, J.M., Doyle, G.V., Allard, W.J., Terstappen, L.W., and Hayes, D.F. (2004). Circulating tumor cells, disease progression, and survival in metastatic breast cancer. N Engl J Med 351, 781–791. 10.1056/NEJMoa040766.

Dahl, K.N., Ribeiro, A.J., and Lammerding, J. (2008). Nuclear shape, mechanics, and mechanotransduction. Circ Res 102, 1307–1318. 10.1161/CIRCRESAHA.108.173989.

De Arcangelis, A., Georges-Labouesse, E., and Adams, J.C. (2004). Expression of fascin-1, the gene encoding the actin-bundling protein fascin-1, during mouse embryogenesis. Gene Expr Patterns 4, 637–643.

Dent, E.W., Gupton, S.L., and Gertler, F.B. (2011). The growth cone cytoskeleton in axon outgrowth and guidance. Cold Spring Harb Perspect Biol 3. 10.1101/cshperspect.a001800.

Desai, R., Sarpal, R., Ishiyama, N., Pellikka, M., Ikura, M., and Tepass, U. (2013). Monomeric alpha-catenin links cadherin to the actin cytoskeleton. Nat Cell Biol 15, 261–273. 10.1038/ncb2685.

Dombroski, J.A., Hope, J.M., Sarna, N.S., and King, M.R. (2021). Channeling the Force: Piezo1 Mechanotransduction in Cancer Metastasis. Cells 10. 10.3390/cells10112815.

Elkhatib, N., Neu, M.B., Zensen, C., Schmoller, K.M., Louvard, D., Bausch, A.R., Betz, T., and Vignjevic, D.M. (2014). Fascin plays a role in stress fiber organization and focal adhesion disassembly. Curr Biol 24, 1492–1499. 10.1016/j.cub.2014.05.023.

Elosegui-Artola, A., Andreu, I., Beedle, A.E.M., Lezamiz, A., Uroz, M., Kosmalska, A.J., Oria, R., Kechagia, J.Z., Rico-Lastres, P., Le Roux, A.L., et al. (2017). Force Triggers YAP Nuclear Entry by Regulating Transport across Nuclear Pores. Cell 171, 1397–1410 e1314. 10.1016/j.cell.2017.10.008.

Farge, E. (2003). Mechanical induction of Twist in the Drosophila foregut/stomodeal primordium. Curr Biol 13, 1365–1377. 10.1016/s0960-9822(03)00576-1.

Fernandez-Sanchez, M.E., Barbier, S., Whitehead, J., Bealle, G., Michel, A., Latorre-Ossa, H., Rey, C., Fouassier, L., Claperon, A., Brulle, L., et al. (2015). Mechanical induction of the tumorigenic beta-catenin pathway by tumour growth pressure. Nature 523, 92–95. 10.1038/nature14329.

Fichtman, B., and Harel, A. (2014). Stress and aging at the nuclear gateway. Mech Ageing Dev 135, 24–32. 10.1016/j.mad.2014.01.003.

Freedman, S.L., Suarez, C., Winkelman, J.D., Kovar, D.R., Voth, G.A., Dinner, A.R., and Hocky, G.M. (2019). Mechanical and kinetic factors drive sorting of F-actin cross-linkers on bundles. Proc Natl Acad Sci U S A 116, 16192–16197. 10.1073/pnas.1820814116.

Friedl, P., Locker, J., Sahai, E., and Segall, J.E. (2012). Classifying collective cancer cell invasion. Nat Cell Biol 14, 777–783. 10.1038/ncb2548.

Friedl, P., and Mayor, R. (2017). Tuning Collective Cell Migration by Cell-Cell Junction Regulation. Cold Spring Harb Perspect Biol 9. 10.1101/cshperspect.a029199.

Fristrom, D. (1988). The cellular basis of epithelial morphogenesis. A review. Tissue Cell 20, 645–690. 10.1016/0040-8166(88)90015-8.

Garcia, M.A., Nelson, W.J., and Chavez, N. (2018). Cell-Cell Junctions Organize Structural and Signaling Networks. Cold Spring Harb Perspect Biol 10. 10.1101/cshperspect.a029181.

George, S.P., Chen, H., Conrad, J.C., and Khurana, S. (2013). Regulation of directional cell migration by membrane-induced actin bundling. J Cell Sci 126, 312–326. 10.1242/jcs.116244.

Grosse-Wilde, A., Fouquier d’Herouel, A., McIntosh, E., Ertaylan, G., Skupin, A., Kuestner, R.E., del Sol, A., Walters, K.A., and Huang, S. (2015). Stemness of the hybrid Epithelial/Mesenchymal State in Breast Cancer and Its Association with Poor Survival. PLoS One 10, e0126522. 10.1371/journal.pone.0126522.

Gunal, A., Onguru, O., Safali, M., and Beyzadeoglu, M. (2008). Fascin expression [corrected] in glial tumors and its prognostic significance in glioblastomas. Neuropathology 28, 382–386. 10.1111/j.1440-1789.2008.00889.x.

Haeger, A., Krause, M., Wolf, K., and Friedl, P. (2014). Cell jamming: collective invasion of mesenchymal tumor cells imposed by tissue confinement. Biochim Biophys Acta 1840, 2386–2395. 10.1016/j.bbagen.2014.03.020.

Hall, E.T., Hoesing, E., Sinkovics, E., and Verheyen, E.M. (2019). Actomyosin contractility modulates Wnt signaling through adherens junction stability. Mol Biol Cell 30, 411–426. 10.1091/mbc.E18-06-0345.

Hashimoto, Y., Skacel, M., Lavery, I.C., Mukherjee, A.L., Casey, G., and Adams, J.C. (2006). Prognostic significance of fascin expression in advanced colorectal cancer: an immunohistochemical study of colorectal adenomas and adenocarcinomas. BMC Cancer 6, 241. 1471-2407-6-241 [pii] 10.1186/1471-2407-6-241.

Hirata, H., Samsonov, M., and Sokabe, M. (2017). Actomyosin contractility provokes contact inhibition in E-cadherin-ligated keratinocytes. Sci Rep 7, 46326. 10.1038/srep46326.

Hoffman, B.D., and Yap, A.S. (2015). Towards a Dynamic Understanding of Cadherin-Based Mechanobiology. Trends Cell Biol 25, 803–814. 10.1016/j.tcb.2015.09.008.

Huang, W., Hu, H., Zhang, Q., Wu, X., Wei, F., Yang, F., Gan, L., Wang, N., Yang, X., and Guo, A.Y. (2019). Regulatory networks in mechanotransduction reveal key genes in promoting cancer cell stemness and proliferation. Oncogene 38, 6818–6834. 10.1038/s41388-019-0925-0.

Hugh, T.J., Dillon, S.A., Taylor, B.A., Pignatelli, M., Poston, G.J., and Kinsella, A.R. (1999). Cadherin-catenin expression in primary colorectal cancer: a survival analysis. Br J Cancer 80, 1046–1051. 10.1038/sj.bjc.6690461.

Humtsoe, J.O., Koya, E., Pham, E., Aramoto, T., Zuo, J., Ishikawa, T., and Kramer, R.H. (2012). Transcriptional profiling identifies upregulated genes following induction of epithelial-mesenchymal transition in squamous carcinoma cells. Exp Cell Res 318, 379–390. 10.1016/j.yexcr.2011.11.011.

Ilina, O., Bakker, G.J., Vasaturo, A., Hofmann, R.M., and Friedl, P. (2011). Two-photon laser-generated microtracks in 3D collagen lattices: principles of MMP-dependent and - independent collective cancer cell invasion. Phys Biol 8, 015010. 10.1088/1478-3975/8/1/015010.

Ilina, O., and Friedl, P. (2009). Mechanisms of collective cell migration at a glance. J Cell Sci 122, 3203–3208. 10.1242/jcs.036525.

Jayo, A., Malboubi, M., Antoku, S., Chang, W., Ortiz-Zapater, E., Groen, C., Pfisterer, K., Tootle, T., Charras, G., Gundersen, G.G., and Parsons, M. (2016). Fascin Regulates Nuclear Movement and Deformation in Migrating Cells. Dev Cell 38, 371–383. 10.1016/j.devcel.2016.07.021.

Jia, D., Jolly, M.K., Tripathi, S.C., Den Hollander, P., Huang, B., Lu, M., Celiktas, M., Ramirez-Pena, E., Ben-Jacob, E., Onuchic, J.N., et al. (2017). Distinguishing mechanisms underlying EMT tristability. Cancer Converg 1, 2. 10.1186/s41236-017-0005-8.

Jolly, M.K., Boareto, M., Huang, B., Jia, D., Lu, M., Ben-Jacob, E., Onuchic, J.N., and Levine, H. (2015). Implications of the Hybrid Epithelial/Mesenchymal Phenotype in Metastasis. Front Oncol 5, 155. 10.3389/fonc.2015.00155.

Jolly, M.K., Tripathi, S.C., Jia, D., Mooney, S.M., Celiktas, M., Hanash, S.M., Mani, S.A., Pienta, K.J., Ben-Jacob, E., and Levine, H. (2016). Stability of the hybrid epithelial/mesenchymal phenotype. Oncotarget 7, 27067–27084. 10.18632/oncotarget.8166.

Joosse, S.A., Gorges, T.M., and Pantel, K. (2015). Biology, detection, and clinical implications of circulating tumor cells. EMBO Mol Med 7, 1–11. 10.15252/emmm.201303698.

Khoo, B.L., Lee, S.C., Kumar, P., Tan, T.Z., Warkiani, M.E., Ow, S.G., Nandi, S., Lim, C.T., and Thiery, J.P. (2015). Short-term expansion of breast circulating cancer cells predicts response to anti-cancer therapy. Oncotarget 6, 15578–15593. 10.18632/oncotarget.3903.

Klymkowsky, M.W., and Savagner, P. (2009). Epithelial-mesenchymal transition: a cancer researcher’s conceptual friend and foe. Am J Pathol 174, 1588–1593. 10.2353/ajpath.2009.080545.

Krendel, M., Gloushankova, N.A., Bonder, E.M., Feder, H.H., Vasiliev, J.M., and Gelfand, I.M. (1999). Myosin-dependent contractile activity of the actin cytoskeleton modulates the spatial organization of cell-cell contacts in cultured epitheliocytes. Proc Natl Acad Sci U S A 96, 9666–9670. 10.1073/pnas.96.17.9666.

Ladoux, B., Nelson, W.J., Yan, J., and Mege, R.M. (2015). The mechanotransduction machinery at work at adherens junctions. Integr Biol (Camb) 7, 1109–1119. 10.1039/c5ib00070j.

le Duc, Q., Shi, Q., Blonk, I., Sonnenberg, A., Wang, N., Leckband, D., and de Rooij, J. (2010). Vinculin potentiates E-cadherin mechanosensing and is recruited to actin-anchored sites within adherens junctions in a myosin II-dependent manner. J Cell Biol 189, 1107–1115. 10.1083/jcb.201001149.

Le, S., Hu, X., Yao, M., Chen, H., Yu, M., Xu, X., Nakazawa, N., Margadant, F.M., Sheetz, M.P., and Yan, J. (2017). Mechanotransmission and Mechanosensing of Human alpha-Actinin 1. Cell Rep 21, 2714–2723. 10.1016/j.celrep.2017.11.040.

Li, A., Morton, J.P., Ma, Y., Karim, S.A., Zhou, Y., Faller, W.J., Woodham, E.F., Morris, H.T., Stevenson, R.P., Juin, A., et al. (2014). Fascin is regulated by slug, promotes progression of pancreatic cancer in mice, and is associated with patient outcomes. Gastroenterology 146, 1386–1396 e1381-1317. 10.1053/j.gastro.2014.01.046.

Li, J., Zhang, S., Pei, M., Wu, L., Liu, Y., Li, H., Lu, J., and Li, X. (2018). FSCN1 Promotes Epithelial-Mesenchymal Transition Through Increasing Snail1 in Ovarian Cancer Cells. Cell Physiol Biochem 49, 1766–1777. 10.1159/000493622.

Li, J.X.H., Tang, V.W., and Brieher, W.M. (2020). Actin protrusions push at apical junctions to maintain E-cadherin adhesion. Proc Natl Acad Sci U S A 117, 432–438. 10.1073/pnas.1908654117.

Lin, S., Taylor, M.D., Singh, P.K., and Yang, S. (2021). How does fascin promote cancer metastasis? FEBS J 288, 1434–1446. 10.1111/febs.15484.

Liu, H., Zhang, Y., Li, L., Cao, J., Guo, Y., Wu, Y., and Gao, W. (2021). Fascin actin-bundling protein 1 in human cancer: promising biomarker or therapeutic target? Mol Ther Oncolytics 20, 240–264. 10.1016/j.omto.2020.12.014.

Ma, Y., Reynolds, L.E., Li, A., Stevenson, R.P., Hodivala-Dilke, K.M., Yamashiro, S., and Machesky, L.M. (2013). Fascin 1 is dispensable for developmental and tumour angiogenesis. Biol Open 2, 1187–1191. 10.1242/bio.20136031.

Mathew, S., George, S.P., Wang, Y., Siddiqui, M.R., Srinivasan, K., Tan, L., and Khurana, S. (2008). Potential molecular mechanism for c-Src kinase-mediated regulation of intestinal cell migration. J Biol Chem 283, 22709–22722. 10.1074/jbc.M801319200.

Michael, M., and Yap, A.S. (2013). The regulation and functional impact of actin assembly at cadherin cell-cell adhesions. Semin Cell Dev Biol 24, 298–307. 10.1016/j.semcdb.2012.12.004.

Na, T.Y., Schecterson, L., Mendonsa, A.M., and Gumbiner, B.M. (2020). The functional activity of E-cadherin controls tumor cell metastasis at multiple steps. Proc Natl Acad Sci U S A 117, 5931–5937. 10.1073/pnas.1918167117.

Nanba, D., Toki, F., Tate, S., Imai, M., Matsushita, N., Shiraishi, K., Sayama, K., Toki, H., Higashiyama, S., and Barrandon, Y. (2015). Cell motion predicts human epidermal stemness. J Cell Biol 209, 305–315. 10.1083/jcb.201409024.

Naphade, S., Sharma, J., Gaide Chevronnay, H.P., Shook, M.A., Yeagy, B.A., Rocca, C.J., Ur, S.N., Lau, A.J., Courtoy, P.J., and Cherqui, S. (2015). Brief reports: Lysosomal cross-correction by hematopoietic stem cell-derived macrophages via tunneling nanotubes. Stem Cells 33, 301–309. 10.1002/stem.1835.

Nieto, M.A., Huang, R.Y., Jackson, R.A., and Thiery, J.P. (2016). Emt: 2016. Cell 166, 21–45. 10.1016/j.cell.2016.06.028.

Niewiadomska, P., Godt, D., and Tepass, U. (1999). DE-Cadherin is required for intercellular motility during Drosophila oogenesis. J Cell Biol 144, 533–547. 10.1083/jcb.144.3.533.

Osswald, M., Jung, E., Sahm, F., Solecki, G., Venkataramani, V., Blaes, J., Weil, S., Horstmann, H., Wiestler, B., Syed, M., et al. (2015). Brain tumour cells interconnect to a functional and resistant network. Nature 528, 93–98. 10.1038/nature16071.

Padmanaban, V., Krol, I., Suhail, Y., Szczerba, B.M., Aceto, N., Bader, J.S., and Ewald, A.J. (2019). E-cadherin is required for metastasis in multiple models of breast cancer. Nature 573, 439–444. 10.1038/s41586-019-1526-3.

Patnaik, S., George, S.P., Pham, E., Roy, S., Singh, K., Mariadason, J.M., and Khurana, S. (2016). By moonlighting in the nucleus, villin regulates epithelial plasticity. Mol Biol Cell 27, 535–548. 10.1091/mbc.E15-06-0453.

Poznansky, M.C., Olszak, I.T., Foxall, R., Evans, R.H., Luster, A.D., and Scadden, D.T. (2000). Active movement of T cells away from a chemokine. Nat Med 6, 543–548. 10.1038/75022.

Przybyla, L., Lakins, J.N., and Weaver, V.M. (2016). Tissue Mechanics Orchestrate Wnt-Dependent Human Embryonic Stem Cell Differentiation. Cell Stem Cell 19, 462–475. 10.1016/j.stem.2016.06.018.

Revenu, C., and Gilmour, D. (2009). EMT 2.0: shaping epithelia through collective migration. Curr Opin Genet Dev 19, 338–342. 10.1016/j.gde.2009.04.007.

Ribeiro, A.S., and Paredes, J. (2014). P-Cadherin Linking Breast Cancer Stem Cells and Invasion: A Promising Marker to Identify an “Intermediate/Metastable” EMT State. Front Oncol 4, 371. 10.3389/fonc.2014.00371.

Ristic, B., Kopel, J., Sherazi, S.A.A., Gupta, S., Sachdeva, S., Bansal, P., Ali, A., Perisetti, A., and Goyal, H. (2021). Emerging Role of Fascin-1 in the Pathogenesis, Diagnosis, and Treatment of the Gastrointestinal Cancers. Cancers (Basel) 13 (*1*), 2536. 10.3390/cancers13112536.

Roper, J.C., Mitrossilis, D., Stirnemann, G., Waharte, F., Brito, I., Fernandez-Sanchez, M.E., Baaden, M., Salamero, J., and Farge, E. (2018). The major beta-catenin/E-cadherin junctional binding site is a primary molecular mechano-transductor of differentiation in vivo. Elife 7, e33381. 10.7554/eLife.33381.

Rotherham, M., and El Haj, A.J. (2015). Remote activation of the Wnt/beta-catenin signalling pathway using functionalised magnetic particles. PLoS One 10, e0121761. 10.1371/journal.pone.0121761.

Rustom, A., Saffrich, R., Markovic, I., Walther, P., and Gerdes, H.H. (2004). Nanotubular highways for intercellular organelle transport. Science 303, 1007–1010.

Scarpa, E., Szabo, A., Bibonne, A., Theveneau, E., Parsons, M., and Mayor, R. (2015). Cadherin Switch during EMT in Neural Crest Cells Leads to Contact Inhibition of Locomotion via Repolarization of Forces. Dev Cell 34, 421–434. 10.1016/j.devcel.2015.06.012.

Schnittler, H., Taha, M., Schnittler, M.O., Taha, A.A., Lindemann, N., and Seebach, J. (2014). Actin filament dynamics and endothelial cell junctions: the Ying and Yang between stabilization and motion. Cell Tissue Res 355, 529–543. 10.1007/s00441-014-1856-2.

Schoumacher, M., El-Marjou, F., Lae, M., Kambou, N., Louvard, D., Robine, S., and Vignjevic, D.M. (2014). Conditional expression of fascin increases tumor progression in a mouse model of intestinal cancer. Eur J Cell Biol 93, 388–395. 10.1016/j.ejcb.2014.08.002.

Seddiki, R., Narayana, G., Strale, P.O., Balcioglu, H.E., Peyret, G., Yao, M., Le, A.P., Teck Lim, C., Yan, J., Ladoux, B., and Mege, R.M. (2018). Force-dependent binding of vinculin to alpha-catenin regulates cell-cell contact stability and collective cell behavior. Mol Biol Cell 29, 380–388. 10.1091/mbc.E17-04-0231.

Shi, S., Zheng, H.C., and Zhang, Z.G. (2020). Roles of Fascin mRNA expression in colorectal cancer: Meta-analysis and bioinformatics analysis. Mol Clin Oncol 13, 119–128. 10.3892/mco.2020.2069.

Simi, A.K., Piotrowski, A.S., Nelson, C.M. . (2015). Mechanotransduction, Metastasis and Genomic Instability (Springer Internation Publishing Switzerland).

Singh, A., and Settleman, J. (2010). EMT, cancer stem cells and drug resistance: an emerging axis of evil in the war on cancer. Oncogene 29, 4741–4751. 10.1038/onc.2010.215.

Smith, H., Whittall, C., Weksler, B., and Middleton, J. (2012). Chemokines stimulate bidirectional migration of human mesenchymal stem cells across bone marrow endothelial cells. Stem Cells Dev 21, 476–486. 10.1089/scd.2011.0025.

Strippoli, A., Cocomazzi, A., Basso, M., Cenci, T., Ricci, R., Pierconti, F., Cassano, A., Fiorentino, V., Barone, C., Bria, E., et al. (2020). c-MYC Expression Is a Possible Keystone in the Colorectal Cancer Resistance to EGFR Inhibitors. Cancers (Basel) 12. 10.3390/cancers12030638.

Taguchi, K., Ishiuchi, T., and Takeichi, M. (2011). Mechanosensitive EPLIN-dependent remodeling of adherens junctions regulates epithelial reshaping. J Cell Biol 194, 643–656. 10.1083/jcb.201104124.

Takeichi, M. (2014). Dynamic contacts: rearranging adherens junctions to drive epithelial remodelling. Nat Rev Mol Cell Biol 15, 397–410. 10.1038/nrm3802.

Tampakis, A., Tampaki, E.C., Nonni, A., Kostakis, I.D., Posabella, A., Kontzoglou, K., von Flue, M., Felekouras, E., Kouraklis, G., and Nikiteas, N. (2021). High fascin-1 expression in colorectal cancer identifies patients at high risk for early disease recurrence and associated mortality. BMC Cancer 21, 153. 10.1186/s12885-021-07842-4.

Tan, V.Y., Lewis, S.J., Adams, J.C., and Martin, R.M. (2013). Association of fascin-1 with mortality, disease progression and metastasis in carcinomas: a systematic review and meta-analysis. BMC Med 11, 52. 10.1186/1741-7015-11-52.

Tebbutt, N.C., Wilson, K., Gebski, V.J., Cummins, M.M., Zannino, D., van Hazel, G.A., Robinson, B., Broad, A., Ganju, V., Ackland, S.P., et al. (2010). Capecitabine, bevacizumab, and mitomycin in first-line treatment of metastatic colorectal cancer: results of the Australasian Gastrointestinal Trials Group Randomized Phase III MAX Study. J Clin Oncol 28, 3191–3198. JCO.2009.27.7723 [pii] 10.1200/JCO.2009.27.7723.

Theveneau, E., Marchant, L., Kuriyama, S., Gull, M., Moepps, B., Parsons, M., and Mayor, R. (2010). Collective chemotaxis requires contact-dependent cell polarity. Dev Cell 19, 39–53. 10.1016/j.devcel.2010.06.012.

Theveneau, E., and Mayor, R. (2011). Can mesenchymal cells undergo collective cell migration? The case of the neural crest. Cell Adh Migr 5, 490–498. 10.4161/cam.5.6.18623.

Thiery, J.P., Acloque, H., Huang, R.Y., and Nieto, M.A. (2009). Epithelial-mesenchymal transitions in development and disease. Cell 139, 871–890. 10.1016/j.cell.2009.11.007.

Ubelmann, F., Chamaillard, M., El-Marjou, F., Simon, A., Netter, J., Vignjevic, D., Nichols, B.L., Quezada-Calvillo, R., Grandjean, T., Louvard, D., et al. (2013). Enterocyte loss of polarity and gut wound healing rely upon the F-actin-severing function of villin. Proc Natl Acad Sci U S A 110, E1380–1389. 1218446110 [pii] 10.1073/pnas.1218446110.

Vianello, F., Olszak, I.T., and Poznansky, M.C. (2005). Fugetaxis: active movement of leukocytes away from a chemokinetic agent. J Mol Med (Berl) 83, 752–763. 10.1007/s00109-005-0675-z.

Vignjevic, D., Schoumacher, M., Gavert, N., Janssen, K.P., Jih, G., Lae, M., Louvard, D., Ben-Ze’ev, A., and Robine, S. (2007). Fascin, a novel target of beta-catenin-TCF signaling, is expressed at the invasive front of human colon cancer. Cancer Res 67, 6844–6853.

Wang, Y., George, S.P., Roy, S., Pham, E., Esmaeilniakooshkghazi, A., and Khurana, S. (2016). Both the anti- and pro-apoptotic functions of villin regulate cell turnover and intestinal homeostasis. Sci Rep 6, 35491. 10.1038/srep35491.

Watanabe, K., Villarreal-Ponce, A., Sun, P., Salmans, M.L., Fallahi, M., Andersen, B., and Dai, X. (2014). Mammary morphogenesis and regeneration require the inhibition of EMT at terminal end buds by Ovol2 transcriptional repressor. Dev Cell 29, 59–74. 10.1016/j.devcel.2014.03.006.

Welch-Reardon, K.M., Ehsan, S.M., Wang, K., Wu, N., Newman, A.C., Romero-Lopez, M., Fong, A.H., George, S.C., Edwards, R.A., and Hughes, C.C. (2014). Angiogenic sprouting is regulated by endothelial cell expression of Slug. J Cell Sci 127, 2017–2028. 10.1242/jcs.143420.

Whitehead, J., Vignjevic, D., Futterer, C., Beaurepaire, E., Robine, S., and Farge, E. (2008). Mechanical factors activate beta-catenin-dependent oncogene expression in APC mouse colon. HFSP J 2, 286–294. 10.2976/1.2955566.

Winkelman, J.D., Suarez, C., Hocky, G.M., Harker, A.J., Morganthaler, A.N., Christensen, J.R., Voth, G.A., Bartles, J.R., and Kovar, D.R. (2016). Fascin- and alpha-Actinin-Bundled Networks Contain Intrinsic Structural Features that Drive Protein Sorting. Curr Biol 26, 2697–2706. 10.1016/j.cub.2016.07.080.

Wolf, C.a.M., M. (2009). “Mechanotransduction: Role of nuclear pore mechanics and nucleocytoplasmic transport,” in Cellular Mechanotransduction: Diverse Perspectives from Molecules to Tissues. (Cambridge University Press, New York).

Wong, M.H., Rubinfeld, B., and Gordon, J.I. (1998). Effects of forced expression of an NH2-terminal truncated beta-Catenin on mouse intestinal epithelial homeostasis. J Cell Biol 141, 765–777. 10.1083/jcb.141.3.765.

Xu, X.M., Yoo, M.H., Carlson, B.A., Gladyshev, V.N., and Hatfield, D.L. (2009). Simultaneous knockdown of the expression of two genes using multiple shRNAs and subsequent knock-in of their expression. Nat Protoc 4, 1338–1348. 10.1038/nprot.2009.145.

Yao, M., Qiu, W., Liu, R., Efremov, A.K., Cong, P., Seddiki, R., Payre, M., Lim, C.T., Ladoux, B., Mege, R.M., and Yan, J. (2014). Force-dependent conformational switch of alpha-catenin controls vinculin binding. Nat Commun 5, 4525. 10.1038/ncomms5525.

Yonemura, S. (2011). Cadherin-actin interactions at adherens junctions. Curr Opin Cell Biol 23, 515–522. 10.1016/j.ceb.2011.07.001.

Yonemura, S., Wada, Y., Watanabe, T., Nagafuchi, A., and Shibata, M. (2010). alpha-Catenin as a tension transducer that induces adherens junction development. Nat Cell Biol 12, 533–542. 10.1038/ncb2055.

Yu, M., Bardia, A., Wittner, B.S., Stott, S.L., Smas, M.E., Ting, D.T., Isakoff, S.J., Ciciliano, J.C., Wells, M.N., Shah, A.M., et al. (2013). Circulating breast tumor cells exhibit dynamic changes in epithelial and mesenchymal composition. Science 339, 580–584. 10.1126/science.1228522.

Zhang, F.R., Tao, L.H., Shen, Z.Y., Lv, Z., Xu, L.Y., and Li, E.M. (2008). Fascin expression in human embryonic, fetal, and normal adult tissue. J Histochem Cytochem 56, 193–199. 10.1369/jhc.7A7353.2007.

Zhang, H.L., Wang, P., Lu, M.Z., Zhang, S.D., and Zheng, L. (2019). c-Myc maintains the self-renewal and chemoresistance properties of colon cancer stem cells. Oncol Lett 17, 4487–4493. 10.3892/ol.2019.10081.

Zheng, Y., Miyamoto, D.T., Wittner, B.S., Sullivan, J.P., Aceto, N., Jordan, N.V., Yu, M., Karabacak, N.M., Comaills, V., Morris, R., et al. (2017). Expression of beta-globin by cancer cells promotes cell survival during blood-borne dissemination. Nat Commun 8, 14344. 10.1038/ncomms14344.

