## Supplementary figures and images for "In colon cancer cells, fascin1 functions as a mechanosensor that transforms adherens junction mechanotransduction"

### Supplementary Figure 1

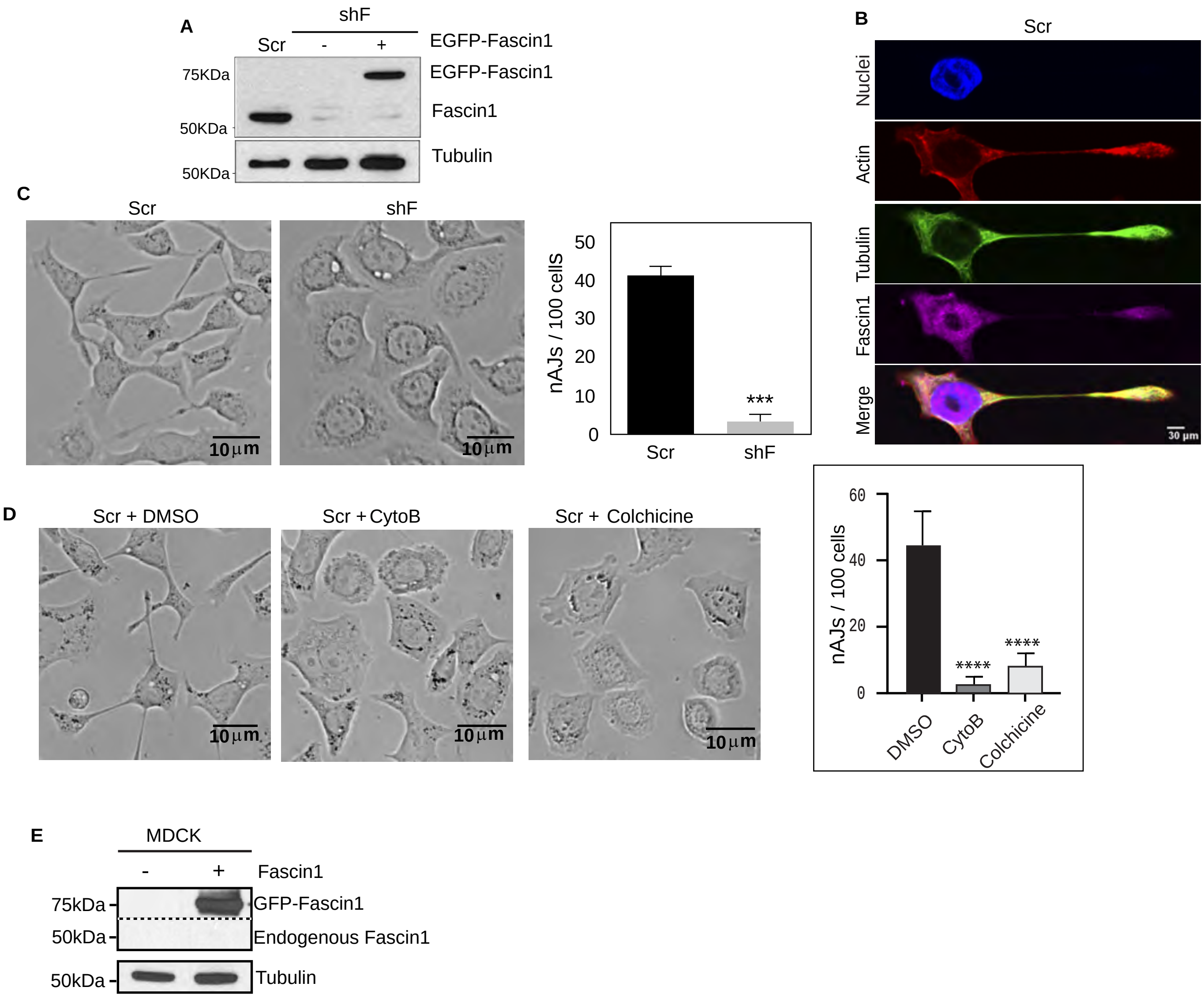

### Supplementary Figure 2

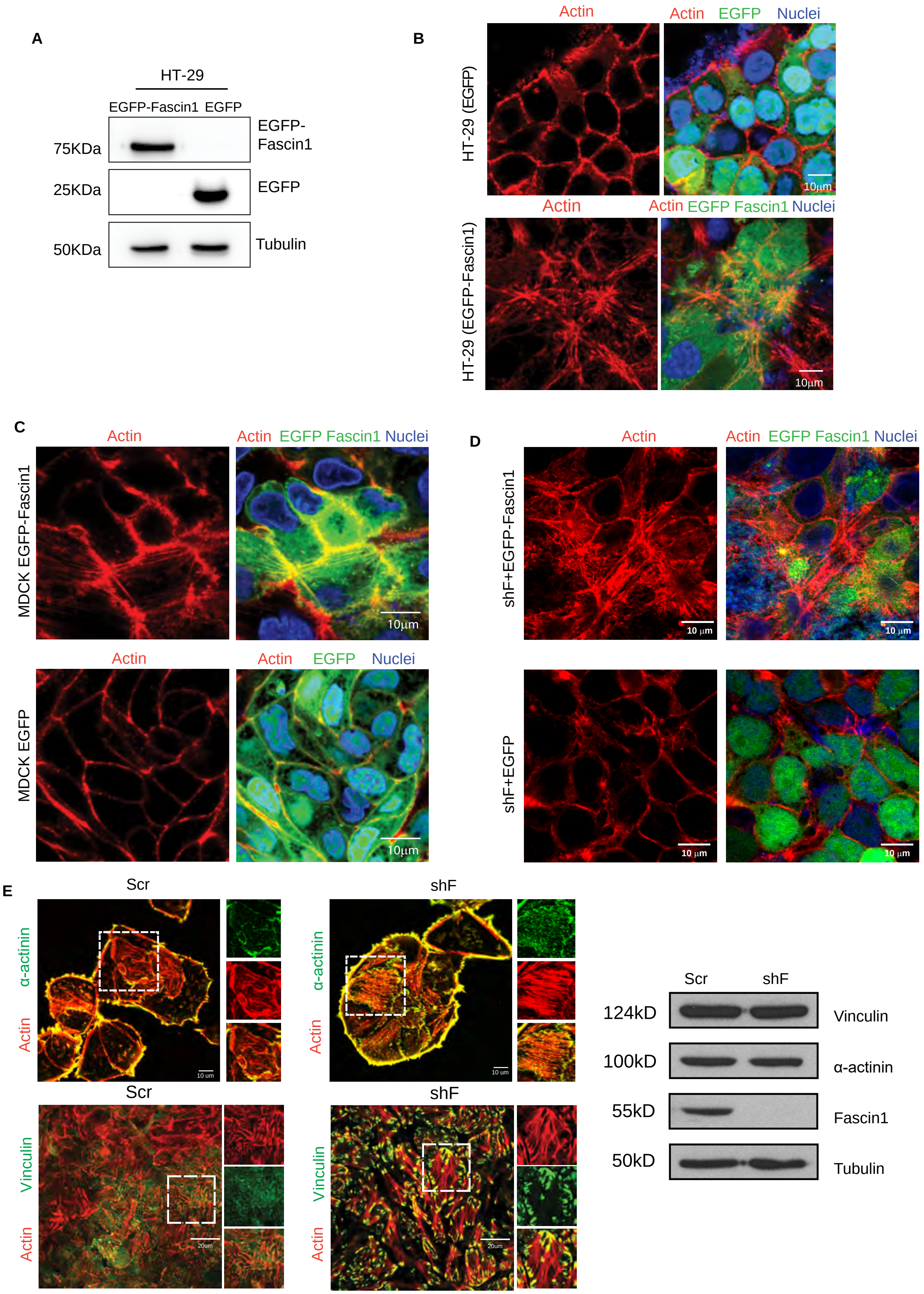

### Supplementary Figure 3

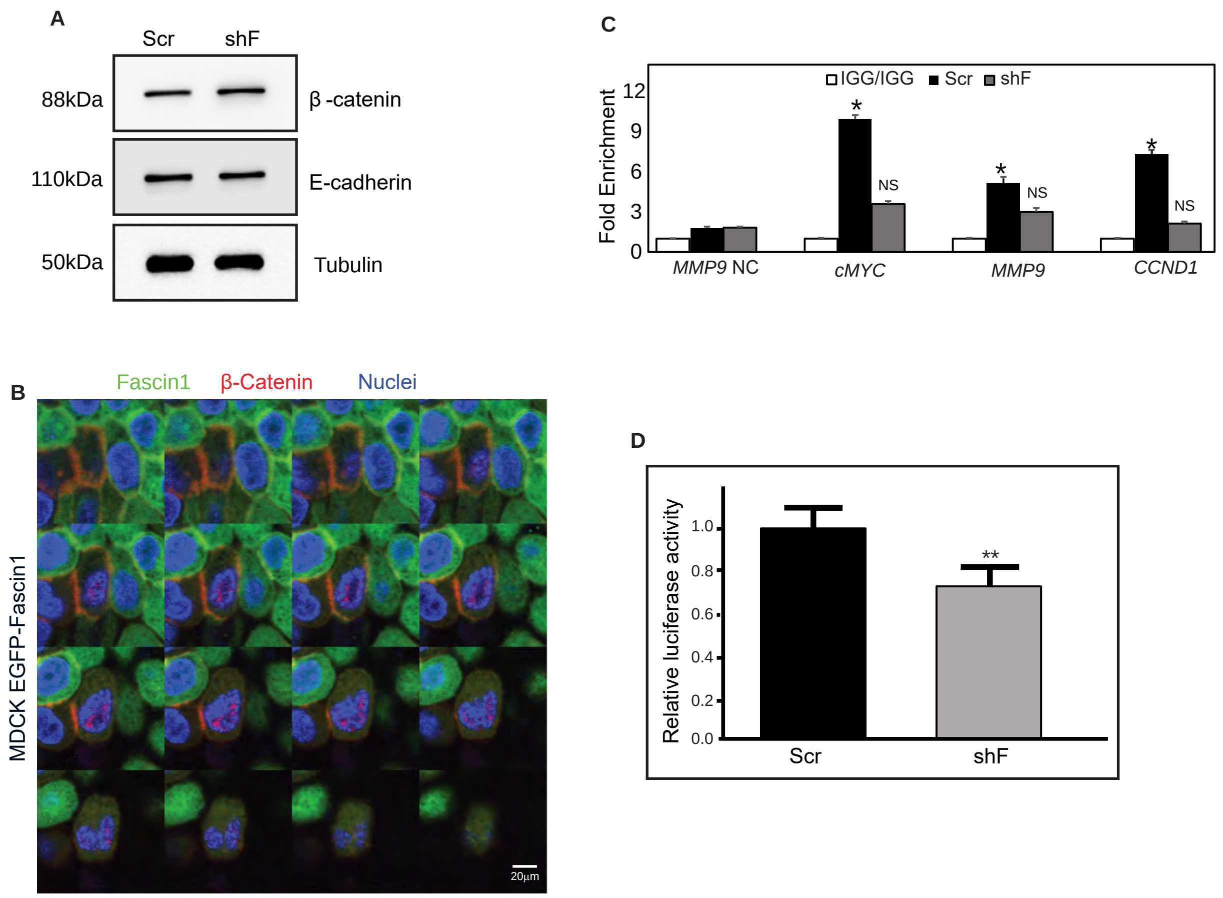

### Supplementary Figure 4

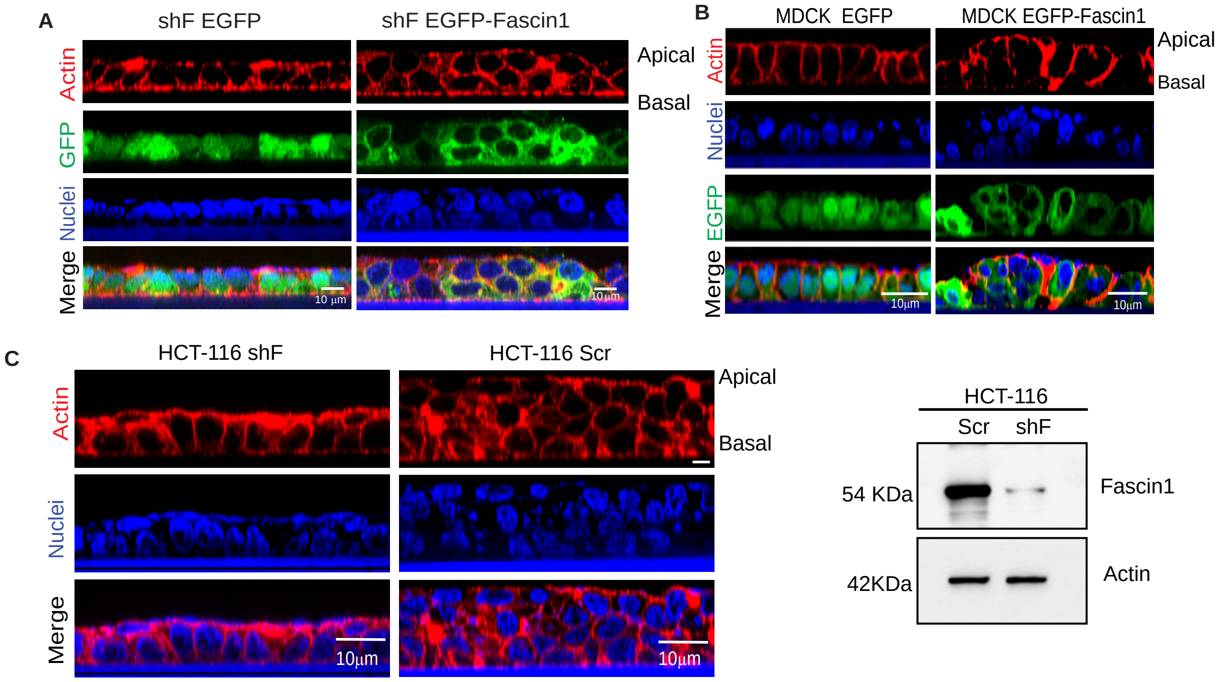

### Supplementary Figure 5

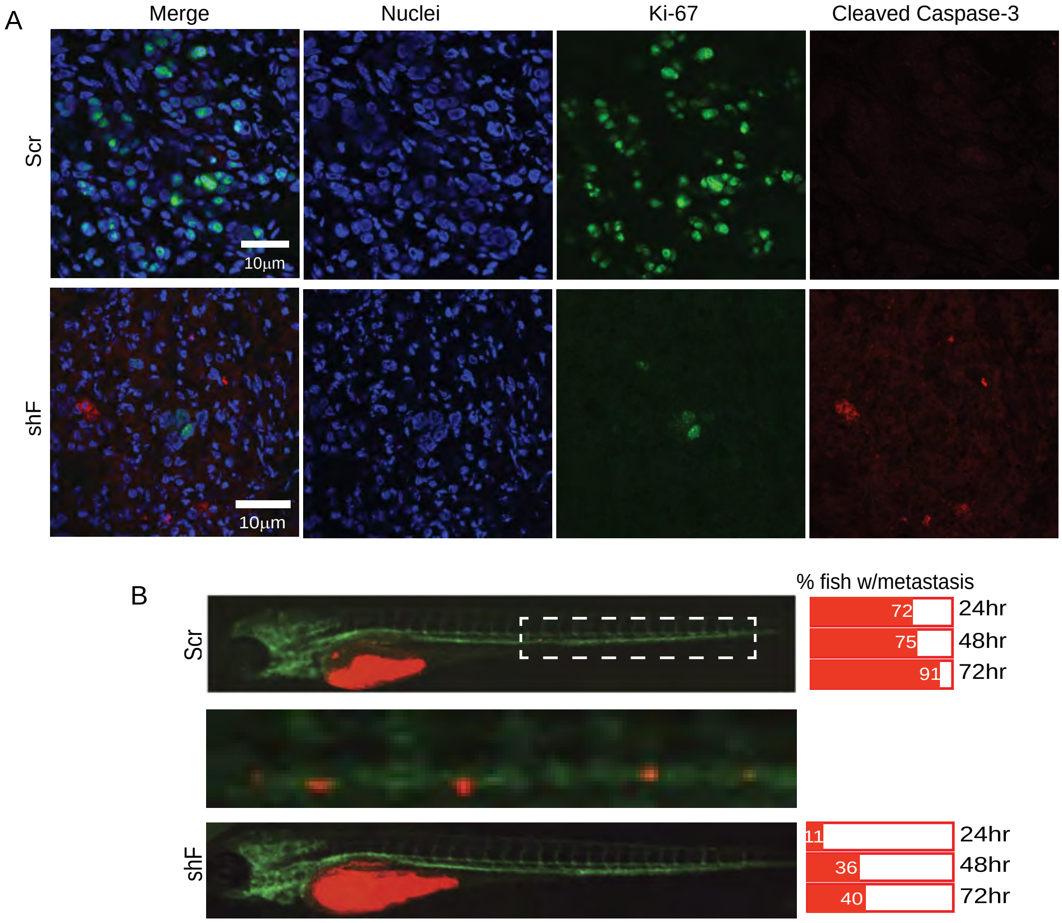

### Supplementary Figure 6

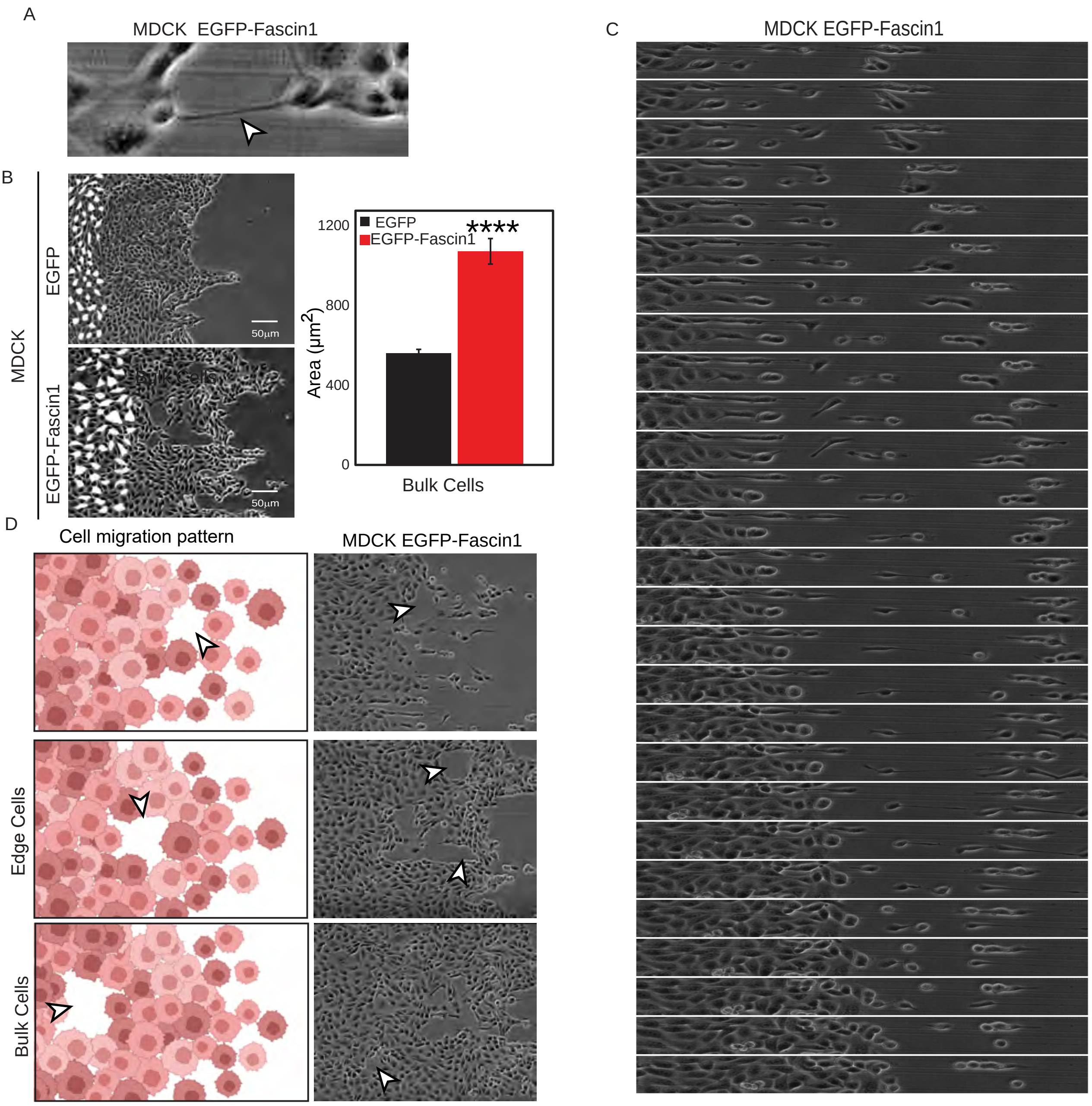
