## Supplementary material for "In colon cancer cells, fascin1 functions as a mechanosensor that transforms adherens junction mechanotransduction": Table

Table 1

Mann-Whitney U Test for elocities, and Kuiper’s Test and Watson U² Test for velocity angles

| Group 1 | Group 2 | Mann-Whitney U Test (p-value) $V_x$ [a] | $V_r$ [b] | $\lvert V \rvert$ [c] | Kuiper’s Test (p-value) θ [d] | Watson $U^2$ (Critical Value) θ |
|---|---|---|---|---|---|---|
| MDCK bulk | MDCK edge | 1.96E-01 | 2.67E-02 | 3.29E-15 | < 0.001 | 11.919* |
| MDCK bulk | EGFP-Fascin1 bulk | 0.00E+00 | 0.00E+00 | 0.00E+00 | < 0.001 | 583.441* |
| MDCK edge | EGFP-Fascin1 edge | 1.46E-51 | 5.46E-52 | 3.75E-36 | < 0.001 | 16.313* |
| EGFP-Fascin1 bulk | EGFP-Fascin1 edge | 7.04E-74 | 3.04E-77 | 9.65E-18 | < 0.001 | 75.251* |
| EGFP-Fascin1 bulk | EGFP-Fascin1 loose | 3.60E-13 | 6.55E-12 | 4.09E-03 | < 0.001 | 160.655* |
| EGFP-Fascin1 edge | EGFP-Fascin1 loose | 1.77E-02 | 5.96E-02 | 3.70E-06 | < 0.001 | 6.605* |
| MDCK all | EGFP-Fascin1 all | 0.00E+00 | 0.00E+00 | 0.00E+00 | | |

[a] $V_x$, X velocity, [b] $V_x$, radial velocity, [c] $\lvert V \rvert$, speed, [d] θ, angle

*For a p-value < 0.05 and n > 100 for sample sets, $U^2$ must be less than 0.1869.
If $U^2$, or the two sample sets are from different distributions.

Table 2

Moments of velocity and velocity angle distributions.

| Group | Parameter | Mean | SD | Skew | Kurtosis |
|---|---|---|---|---|---|
| MDCK bulk | $V_x$ | 0.73 µm/min | 0.97 µm/min | -0.27 | 7.82 |
| MDCK edge | $V_x$ | 0.69 µm/min | 1.32 µm/min | -1.26 | 10.08 |
| EGFP-Fascin1 bulk | $V_x$ | 1.43 µm/min | 1.00 µm/min | -1.73 | 15.80 |
| EGFP-Fascin1edge | $V_x$ | 1.09 µm/min | 1.38 µm/min | -0.84 | 6.55 |
| EGFP-Fascin1 loose | $V_x$ | 0.82 µm/min | 1.90 µm/min | -0.40 | 1.04 |
| MDCK bulk | $V_r$ | 0.80 µm/min | 0.94 µm/min | 0.01 | 8.05 |
| MDCK edge | $V_r$ | 0.75 µm/min | 1.23 µm/min | -0.38 | 9.06 |
| EGFP-Fascin1bulk | $V_r$ | 1.45 µm/min | 0.99 µm/min | -1.57 | 16.01 |
| EGFP-Fascin1edge | $V_r$ | 1.13 µm/min | 1.34 µm/min | -0.52 | 6.00 |
| EGFP-Fascin1 loose | $V_r$ | 0.98 µm/min | 1.76 µm/min | 0.15 | 0.99 |
| MDCK bulk | θ | 0.57 θ =13.35 | 0.65 | 0.19 θ =58.99 | 0.14 θ =-122.02 |
| MDCK edge | θ | 0.51 θ =8.67 | 0.70 | 0.12 θ =148.16 | 0.20 θ =-29.52 |
| EGFP-Fascin1 bulk | θ | 0.85 θ =1.95 | 0.39 | 0.47 θ =-150.16 | 2.31 θ =15.58 |
| EGFP-Fascin1 edge | θ | 0.65 θ =-0.11 | 0.59 | 0.72 θ =-176.30 | 1.75 θ =6.48 |
| EGFP-Fascin1 loose | θ | 0.40 θ =8.37 | 0.77 | 0.58 θ =-152.92 | 0.94 θ =22.84 |

[a] $V_x$, X velocity, [b] $V_x$, radial velocity, [c] $\lvert V \rvert$, speed, [d] θ, angle
